# Structural principles of transcriptional collisions

**DOI:** 10.64898/2026.04.06.716678

**Authors:** John W. Watters, Andreas U. Mueller, Xiangwu Ju, Stephany J. Chuquimarca, Hannah J. Ye, Seth A. Darst, Gregory M. Alushin, Shixin Liu

## Abstract

RNA polymerase (RNAP) must navigate crowded genomic tracks to robustly produce transcripts. DNA-bound proteins act as transcription-impeding roadblocks, which are eventually overcome by RNAPs through unclear mechanisms. Here, we use cryo-electron microscopy to visualize actively transcribing *E. coli* RNAP upon collision with an inactivated restriction enzyme (EcoRI*) or with another converging RNAP. Both collisions induce RNAP backtracking into an inactive swiveled state. Swiveling is coupled to DNA deformation through a characteristic structural landscape, mediating RNAP pause stabilization across distinct collision geometries. EcoRI* roadblock bypass efficiency is impacted by factors that modulate RNAP swiveling, backtracking rescue, and roadblock stability. In comparison, head-on RNAP-RNAP collisions feature substantial heterogeneity suggestive of sustained dynamics, with variable inter-RNAP distances that are modulated by nascent transcript hairpins which position bidirectional termination sites. By resolving, to our knowledge, novel structures of actively transcribing RNAP undergoing collisions, we provide a mechanistic framework for interpreting mechanical conflicts during transcription.

## Introduction

The genome has been described as a molecular highway where the macromolecular machines that carry out fundamental central dogma processes (e.g. replication and transcription) function in a densely crowded environment^1^. This milieu includes a variety of static roadblocks (e.g. DNA-binding proteins and DNA lesions), signals (regulatory sequences), and other dynamic machines operating on the same substrate, such that collisions *in vivo* are inevitable^1,2^. In addition to interfering with machine function^2^, collisions also provide potential for regulation, for example, by defining genomic boundaries or coupling the activities of multiple machines in tandem^3,4^, thereby enhancing biological robustness.

Recent studies have emphasized that multiple categories of collisions between DNA machines can have both positive and negative biological consequences. For example, head-on collisions between replication and transcription machinery lead to increased R-loop formation, which is detrimental to genome stability^5–7^. Conversely, local chromatin looping into topologically associated domains is driven by cohesin-CTCF collisions, mediating functional genome organization^8^. Furthermore, co-directional collisions between machines can enhance their processivity, such as translation-transcription coupling in bacteria, which prevents transcriptional arrest^9–15^. Additionally, we recently showed that sequence-specific pausing of bacterial RNA polymerases (RNAPs) positions RNAP-RNAP head-on collisions, thereby programing transcription termination sites to enhance the production of precise transcripts from convergent genes^16,17^. How collisions both disrupt and enhance machine functions at the protein structural level to produce these diverse outcomes remains unknown.

Because of its central role in many genomic collisions, here we focus on the RNAP as a model machine, which operates as a Brownian rachet and can generate strong forces and torques^18–23^. Previous biophysical studies of RNAP collisions have suggested that upon being halted by roadblocks, the complex can bypass them by two mechanisms: (i) passively waiting for roadblocks to dissociate or (ii) actively consuming chemical energy to dislodge them^24^, with the relative contributions of these mechanisms dependent on roadblock identity^15,16,24,25^. Additional factors including transcription elongation factors, co-directional transcribing RNAPs, and applied force have also been shown to help RNAP overcome roadblocks^15,16,24^. While RNAP has been extensively characterized by single-molecule biophysics, structural biology, and biochemistry, it remains unclear how roadblocks effectively stall actively transcribing RNAPs and how RNAPs in turn bypass roadblocks. Collisions occur under non-equilibrium, actively energy-dissipating conditions that are technically challenging to recapitulate for structural studies. Conceptually, it is interesting to examine how collisions involving a nanoscale machine like RNAP—which operates under the low Reynolds number regime—compared to macroscopic collisions such as a car crash, which produces heterogeneous outcomes in the final motor configuration. However, such microscopic transcriptional collisions have rarely been directly visualized at the atomic level.

In this study, we use cryo-electron microscopy (cryo-EM) to examine actively transcribing *E. coli* RNAPs collided with (i) a static roadblock, namely the catalytically deficient EcoRI E112Q restriction enzyme mutant (hereafter referred to as EcoRI*), and (ii) another RNAP transcribing in the opposite direction, resulting in a head-on collision. By comparing the structural landscapes of these two physiological collision scenarios^24,26,27^, we uncover principles for mechanical control of RNAP that can be harnessed to bypass deleterious roadblocks and mediate functional positioning of transcription termination sites.

## Results

### Reconstituting RNAP collision with a stationary roadblock

To capture a collided roadblock complex, we generated a nucleic acid scaffold containing a pre-formed transcription bubble and loaded the scaffold with an RNAP core enzyme and a downstream EcoRI* dimer bound to the cognate binding site (5’-GAATTC-3’, *K*_D_ ∼2.5 fM)^26,27^. Upon ribonucleotide (NTP) addition, RNAP is anticipated to transcribe for ∼20 nucleotides (nt) before encountering the EcoRI* roadblock (**Fig. 1A**). Proper loading of RNAP and EcoRI* was confirmed via a native gel-shift assay, and the complex was enriched using gel filtration chromatography (**Fig. 1B, S1**). We used a Cy3-labeled RNA in the scaffold to evaluate the length of transcription products. In the absence of EcoRI*, most RNAPs transcribed until the end of the DNA template, generating run-off transcripts ∼70 nt in length (**Fig. 1C**). When EcoRI* was added, shorter RNAs (narrowly distributed around ∼40 nt in length) were observed with minimal run-off products (**Fig. 1C**), suggesting high occupancy of EcoRI* and efficient blockage of RNAP readthrough.

**Figure 1.**
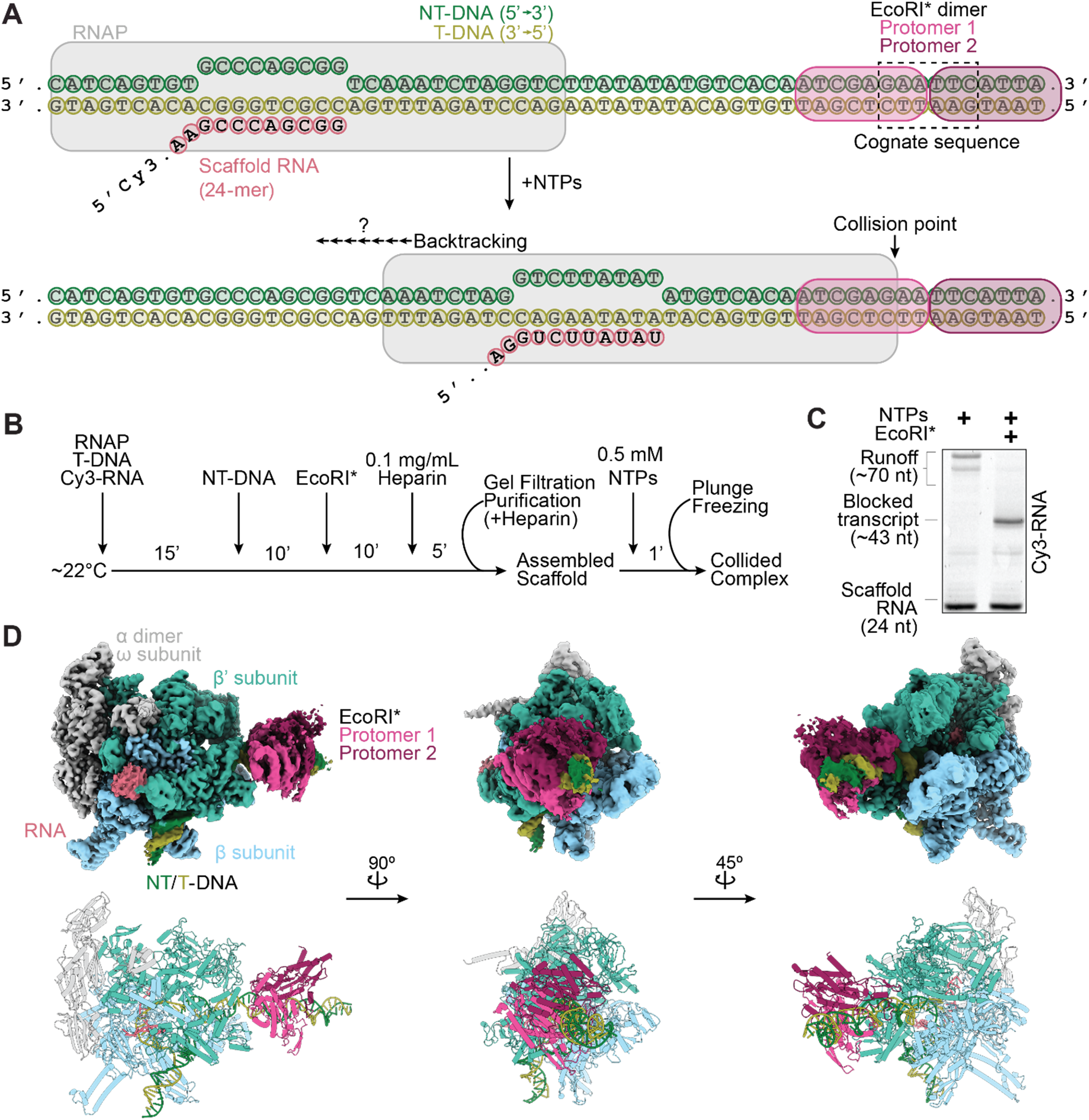
Reconstituting transcriptional collision with a stationary roadblock. **[A]** Synthetic nucleic acid scaffold design to assemble E. coli RNAP and EcoRI* onto a DNA template. NT-DNA, non-template DNA strand; T-DNA, template DNA strand. **[B]** Scaffold assembly and purification scheme. **[C]** Representative denaturing gel showing blocked transcription in the presence of EcoRI*, suggesting collided complex assembly. Cy3-labeled RNA was visualized. **[D]** Cryo-EM density map (top) and atomic model (bottom) of the RNAP-EcoRI* collided complex.

We next used the RNAP-EcoRI* assembly supplemented with 1.5 mM Fos-Choline-8 detergent for cryo-EM sample preparation. To induce collision, 0.5 mM NTPs were added and incubated for 60 seconds prior to plunge-freezing (**Fig. 1B**). After data collection, we performed 3D classification to enrich for particles that contained both RNAP and EcoRI*. Unexpectedly, we obtained a single three-dimensional (3D) reconstruction of the RNAP-EcoRI* collided complex with a global Fourier shell correlation (FSC) resolution of 2.68 Å, indicating that we captured a single metastable state instead of multiple states along the collision pathway (**Fig. S2, S3A-D**). The resolution of the downstream EcoRI* roadblock was significantly lower than the RNAP core, suggesting higher flexibility^26,28,29^. We used 3DFlex in cryoSPARC to improve the resolution of the flexible downstream DNA and EcoRI*, producing a reconstruction with a global resolution of 2.83 Å that allowed us to build an atomic model of the entire collided complex (**Fig. 1D, S3E-H, Table S2**)^30^.

### EcoRI*-collided RNAP is backtracked

We envisioned three possible scenarios for the behavior of RNAP upon colliding with a roadblock: (1) it could advance to the final position along the DNA template free from steric hindrance and remain there; (2) it could halt before this final position due to topological constraints imposed by the DNA; or (3) it could recoil away from the final position to an energetically more favorable site, due to the ability of RNAP to backtrack along DNA^31,32^, as suggested by previous studies of RNAP transcription in the presence of an EcoRI* roadblock^15,24^. To assess RNAP backtracking in our system, we took advantage of the fact that backtracked RNA is sensitive to GreA/B-stimulated endonucleolytic cleavage^33^. In the absence of GreA/B, the majority of RNA products were 43 nt long (**Fig. 2A),** while in the presence of GreB and, to a lesser extent, GreA, the distribution of bands was shifted towards shorter lengths. In the presence of both factors, most RNA products were 39-41 nt in length (**Fig. 2A, S4A, B**), suggesting that the collided RNAP is backtracked 2-4 nt in our reconstitution system, in agreement with previous results^15^.

**Figure 2.**
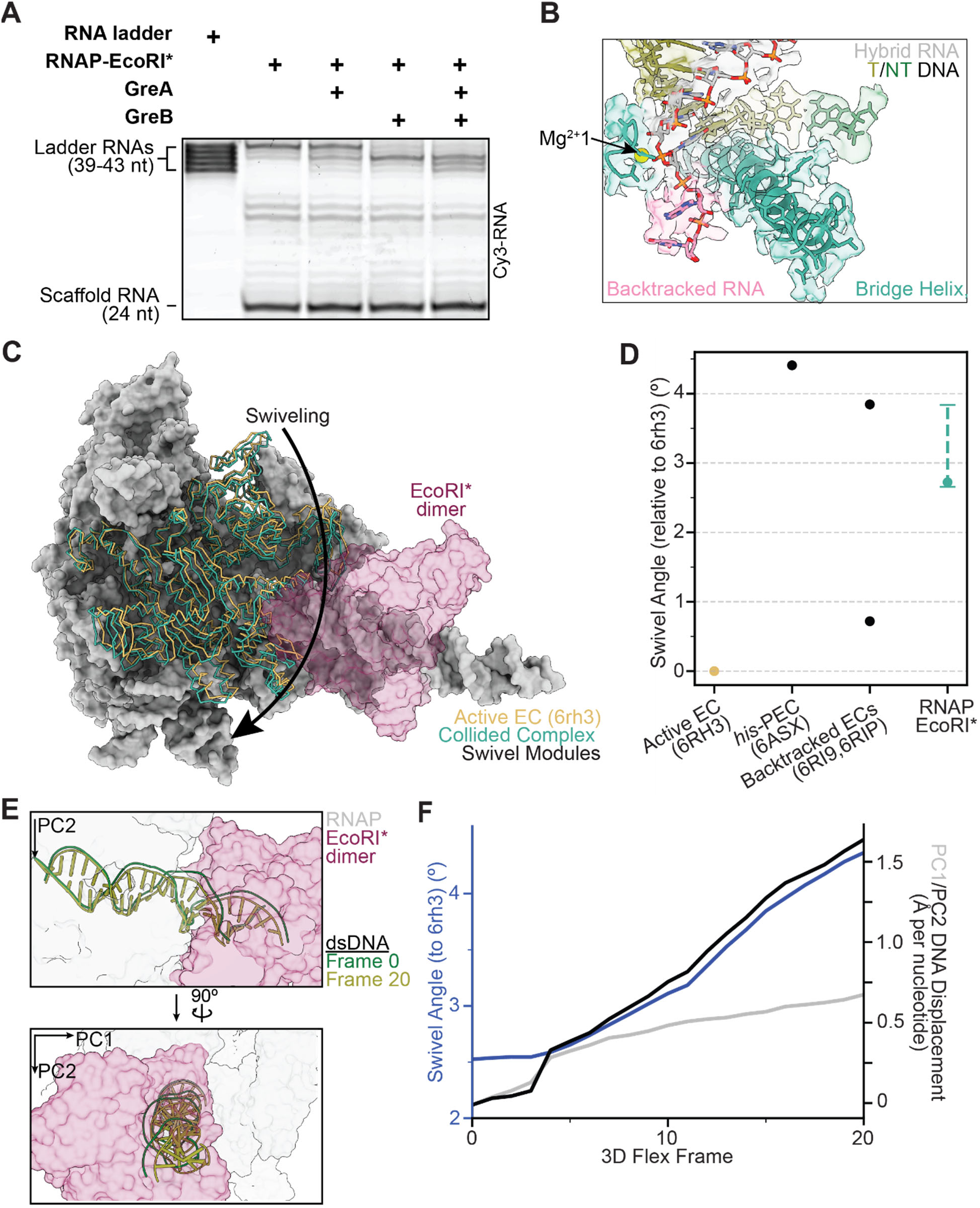
EcoRI* collided complex is backtracked and moderately swiveled. [A]. Representative gel of in vitro transcription assay in the presence and absence of Gre factors, with an RNA ladder corresponding to potential transcript lengths within the collided complex. **[B]** View of the active site of the collided complex (cryo-EM map and atomic model). Resolvable backtracked bases are highlighted in pink. **[C]** View of the swivel module of an active elongation complex (PDB: 6RH3) versus that of the EcoRI* collided complex, superimposed on active EC structure after alignment, highlighting differential swiveling. **[D]** Swivel angle magnitudes of prior RNAP structures versus that of the RNAP-EcoRI* collided complex. The value for the consensus reconstruction is displayed as a solid dot, while the range measured across all five 3D flex trajectories for the collided complex is displayed as a dashed line. **[E]** Initial and final frames of the trajectory displayed in **F**, highlighting DNA displacement. **[F]** Swivel angle and DNA displacement measured for each frame of the first series (accounting for the largest conformational movements) of the 3DFlex variability trajectory for the collided complex.

During each nucleotide addition cycle, conformational changes in the bridge helix and trigger loop—structural elements in the RNAP active site—are coordinated to translocate the DNA register, facilitate nucleotide binding, and accelerate the catalysis of nucleotide incorporation^34^. We next examined the RNAP active site in our cryo-EM map and found that it resembles previously reported structures of backtracked RNAP (PDB: 6RI9, 6RIP), featuring an unfolded trigger loop and a straight bridge helix (**Fig. 2B**)^35,36^. Moreover, we observed densities in the secondary channel consistent with extruded, backtracked RNA (**Fig. 2B**). Two backtracked nucleotides are resolved; the lack of density from additional backtracked nucleotides is likely attributable to higher levels of conformational flexibility, as has been observed in previous cryo-EM structures of backtracked RNAP^35^.

These biochemical and structural data suggest that the RNAP transcribes 19 nt of RNA from its initial position, but it cannot stably reside at that position, potentially due to steric clashes with the roadblock. After collision, RNAP backtracks by as many as 4 nt, allowing it to re-equilibrate at a more favorable position. To further evaluate this hypothesis, we computationally modeled the collision pathway by moving RNAP structures to each nucleotide position along the collision axis (up to 44 nt, one nt past the final position transcribed by RNAP in the EcoRI* collision scenario). We then measured the overlapping volume enclosed by the solvent-excluded surfaces (SES) of RNAP and EcoRI* as a proxy of steric overlap (**Fig. S4C, D**). The RNAP conformation was modeled based on a previously reported elongation complex (EC) structure (PDB: 6RH3)^35^. When the polymerase was placed less than two bases away from the collision interface, the overlapping SES volume^37^ increases substantially, consistent with clashes occurring. While this analysis is limited by the assumption that the RNAP retains the same conformation as it approaches the roadblock, it nonetheless indicates that direct steric hindrance from EcoRI* is a plausible explanation for why RNAP backtracks away from the collision site.

### EcoRI*-collided RNAP is swiveled

We next examined the RNAP structural modules that influence its catalytic activity. We find that in the collided RNAP-EcoRI* complex, RNAP is in a ‘swiveled’ conformation, in which a swivel module (β subunit: AA 1241-1341; β’ subunit: AA 1-342, 369-420, 787-930, 1135-1375) is rotated relative to the RNAP structural core. When RNAP is sufficiently swiveled, the bridge helix kinks and the trigger loop is unable to fold into the correct conformation to mediate catalysis, stabilizing long-lived transcriptional pausing^38–40^. In the active elongation complex (EC; PDB: 6rh3)^35^, RNAP is in a non-swiveled state, whereas pronounced swiveling (∼4.5-6° swivel module rotation relative to the active EC) is observed in various paused and backtracked ECs^35,36,39,41^.

In the collided RNAP-EcoRI* complex, the RNAP adopts a moderately swiveled state with a swivel module rotation of 2.8° relative to the active EC, a lower magnitude rotation than observed in a previously reported hairpin-stabilized paused EC (4.4°, PDB: 6ASX)^39^ (**Fig. 2C, D**). We then analyzed 3DFlex variability trajectories to assess swiveling heterogeneity in our data. We found that the swivel angle ranged between 2.6° and 3.8° across frames (**Fig. 2D**)^30^. In contrast, previously resolved backtracked RNAPs populate either swiveled (3.8°, 63% of population, PDB: 6RIP) or non-swiveled states (0.7°, 37% of population, PDB: 6RI9), but not in between (**Fig. 2D**)^35^. This result indicates that the collided RNAP is constrained within a moderately swiveled state, suggesting a novel configuration imposed by the physical collision.

EcoRI binding to DNA by itself is known to distort DNA from its canonical B-form at the cognate sequence, introducing multiple kinks that result in a net 25° bending of the DNA and concomitant increased accessibility of bases in the major groove^28,42^. Given the ability of DNA to serve as a mechanical transducer, we hypothesized that DNA distortions could be transmitted to RNAP as a mechanism to indicate the presence of a roadblock. To investigate coupling between DNA deformation and RNAP structural remodeling, we once again analyzed 3DFlex trajectories. To assess changes in DNA positioning relative to the active EC (PDB: 6RH3), a reasonable proxy for the spatial orientation of DNA relative to the RNAP structural core during catalysis, we established a frame of reference by subjecting the downstream nucleotides to principal component analysis (PCA). The displacement of downstream DNA in the RNAP-EcoRI* complex was then measured from projections onto the PC1 and PC2 orthoplanes after superimposing the structural cores of the RNAPs. Interestingly, we observed that specific DNA distortions downstream from the RNAP active site are correlated with changes in the RNAP swiveling angle (**Fig. 2E, S5, Movie S1**). As the downstream DNA is displaced more along PC2, the magnitude of RNAP swiveling increases (**Fig. 2F, S5**). This observation suggests that modulation of DNA substrate geometry can couple inhibitory RNAP rearrangements to the presence of roadblocks at collision sites.

### Factors that modulate RNAP backtracking / swiveling impact bypass efficiency

Because RNAP is backtracked and swiveled in the RNAP-EcoRI* complex, we next examined whether the outcome of this collision (persistent RNAP halting versus roadblock bypass) can be biased by factors that modulate these aspects of the RNAP structural landscape. We implemented a bulk biochemical assay to quantitatively evaluate roadblocking efficiency (*E*_RB_), conducting transcription reactions at 37 °C for 30 minutes with increasing salt concentrations (50 to 500 mM KOAc) to permit measurable levels of bypass. As anticipated, higher levels of roadblock bypass (lower *E*_RB_) were observed at higher salt concentrations, presumably due to destabilization of the EcoRI*-DNA interaction (**Fig. 3A, B**). Addition of NusA (a pro-swiveling factor) and NusG (an anti-swiveling, anti-backtracking factor)^36,43,44^ led to decreased and increased bypass, respectively (normalized *E*_RB_ = 1.2 ± 0.1 for NusA and 0.8 ± 0.1 for NusG in the presence of EcoRI*, compared to 1.0 ± 0.1 for EcoRI* alone) (**Fig. 3C**). These results suggest that biochemically modulating RNAP’s swiveling propensity (e.g. through Nus factor engagement) changes its capacity to overcome roadblocks. Strikingly, the presence of the transcript cleavage factor GreB, which rescues backtracked RNAP^33^, dramatically reduced roadblocking efficiency (normalized *E*_RB_ = 0.06 ± 0.03) (**Fig. 3B, C**). This near-complete roadblock bypass can be explained by repeated collisions between RNAP and EcoRI* made possible by GreB-induced transcription restart after RNAP backtracking, consistent with a recently proposed ‘battering’ model^24^.

**Figure 3.**
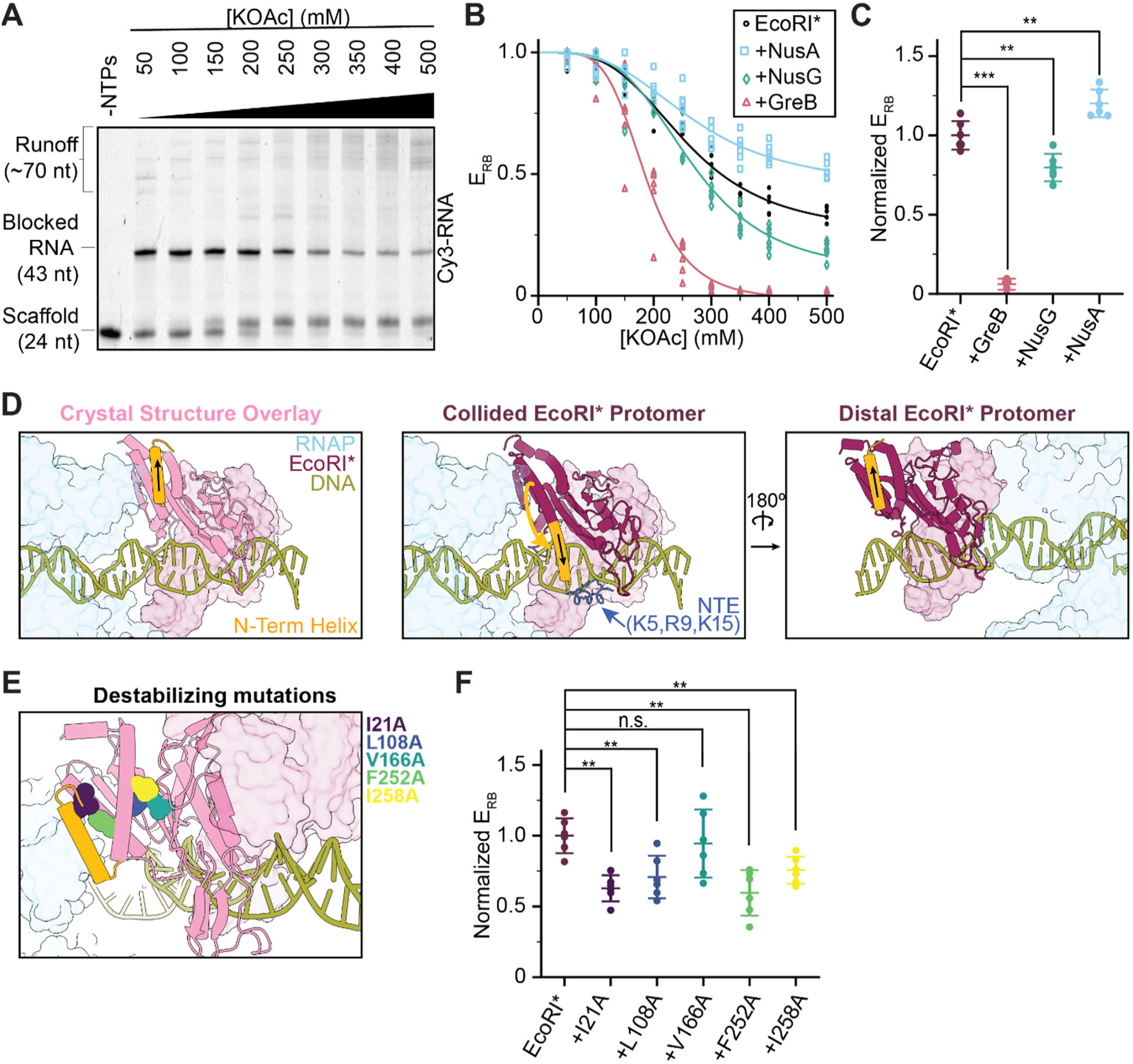
Factors that affect bypass efficiency. **[A]** Representative EcoRI* bypass assay. The RNA lengths under different conditions are visualized via Cy3 fluorescence. **[B]** Fraction of blocked RNAP in the absence and presence of factors (GreB, NusA, NusG). Raw measurements and their sigmoidal fit for each condition are displayed. **[C]** Normalized roadblock efficiencies in the presence of indicated factors at 300 mM KOAc. Data are shown as mean ± S.D. Individual points are also displayed (N = 6). **[D]** Views highlighting the orientation of EcoRI* N-terminal helix in the collided complex versus in the EcoRI* crystal structure (PDB 1ERI). The helix of the collided EcoRI* protomer is flipped and re-oriented toward the downstream dsDNA, which may be stabilized by NTE-DNA interactions, while the distal, non-collided protomer (right) adopts a conformation similar to the crystal structure. **[E]** Positions of single alanine mutations are highlighted on the crystal structure overlay. **[F]** Normalized roadblock efficiencies for the destabilizing EcoRI* mutations at 300 mM KOAc. Data are shown as mean ± S.D. Individual points are also displayed (N = 6). Conditions were compared via two-tailed, unpaired t test including Welch’s correction and Benjamini-Hochberg correction for multiple testing. n.s., not significant; *p < 0.05; **p < 0.01; ***p < 0.001.

### Roadblock structural remodeling facilitates bypass

Having observed substantial structural alterations in RNAP upon collision, we next examined whether the structure of the EcoRI* roadblock is also impacted. EcoRI* engages DNA as a homodimer^28,42,45^. The protein’s DNA-bound conformation is overall highly similar between the RNAP-EcoRI* collided complex and a previous crystal structure in the absence of RNAP (PDB: 1ERI)^42^, with the notable exception for the N-terminal helix (AA 20-31) of the protomer proximal to the collision interface, which flips ∼150°, orienting it away from RNAP (**Fig. 3D, Movie S2**). This rearrangement was not observed in the distal EcoRI* protomer, suggesting it is specifically linked to contact with RNAP (**Fig. 3D**). We speculate that it occurs at the collision site to reduce steric clashes with RNAP, becoming kinetically trapped as the RNAP backtracks away. Additionally, this rearrangement repositions EcoRI*’s N-terminal extension (NTE) residues (5K, 9R, 15K) in an orientation where they can access DNA (**Fig. 3D**). These amino acids have been previously shown to enhance DNA binding in biochemical assays, presumably by mediating electrostatic interactions^26,28,42,45^.

In the EcoRI* crystal structure, the N-terminal helix forms hydrophobic contacts with the protein’s structural core^42^. As a protein’s potency as a roadblock intuitively depends on its mechanical resilience, we posited that destabilizing the overall folded structure of EcoRI* would compromise its roadblocking efficacy. To test this hypothesis, we introduced alanine substitutions at the positions of hydrophobic residues within the EcoRI* core (**Fig. 3E**), distal from the DNA binding interface, and assessed their impact on *E*_RB_ using our roadblock assay. We found that all point mutations (with the exception of V166A) led to significantly increased RNAP bypass (decreased *E*_RB_) (**Fig. 3F**). Notably, the two mutations that caused the largest magnitude changes, I21A and F252A, are internally facing residues adjacent to the predicted collision interface that are likely to mediate a stabilizing contact between the N-terminal helix and the EcoRI* core (**Fig. 3E, F**). These results suggest that the structural stability of a roadblock protein—independent from its affinity to DNA—is linked to its capacity to resist physical impacts from motors like RNAP; correspondingly, mechanical destabilization of roadblocks can enhance bypass.

### Reconstituting head-on collisions between two RNAPs

To explore the generality of RNAP’s collision response, we next investigated another physiological scenario, when an RNAP runs into another active motor complex rather than a stationary roadblock. In previous studies, we demonstrated that head-on collisions between two RNAPs produce long-lived complexes on DNA^16^, constituting a prevalent mechanism for transcription termination of convergent genes in *E. coli*^17^. To reconstitute a head-on RNAP-RNAP collision in a format suitable for cryo-EM structural interrogation, we constructed a DNA template with two convergent promoters (**Fig. 4A**). Each promoter was bound by a σ^70^-RNAP holoenzyme that was walked to the +18/+20 position by omitting one of the four nucleotides (CTP), followed by purification via size exclusion chromatography. Transcription was then restarted with addition of 1 mM NTPs to induce a collision (**Fig. 4B**). We also introduced a native sequence from the *E. coli* genomic region between the *YnaJ*-*UspE* gene pair into the template, which encodes an RNA hairpin containing an 8-bp stem. We have previously shown that this sequence is important for positioning the collision site between a pair of RNAPs, mediating precise termination of convergent transcription^16,17^.

**Figure 4.**
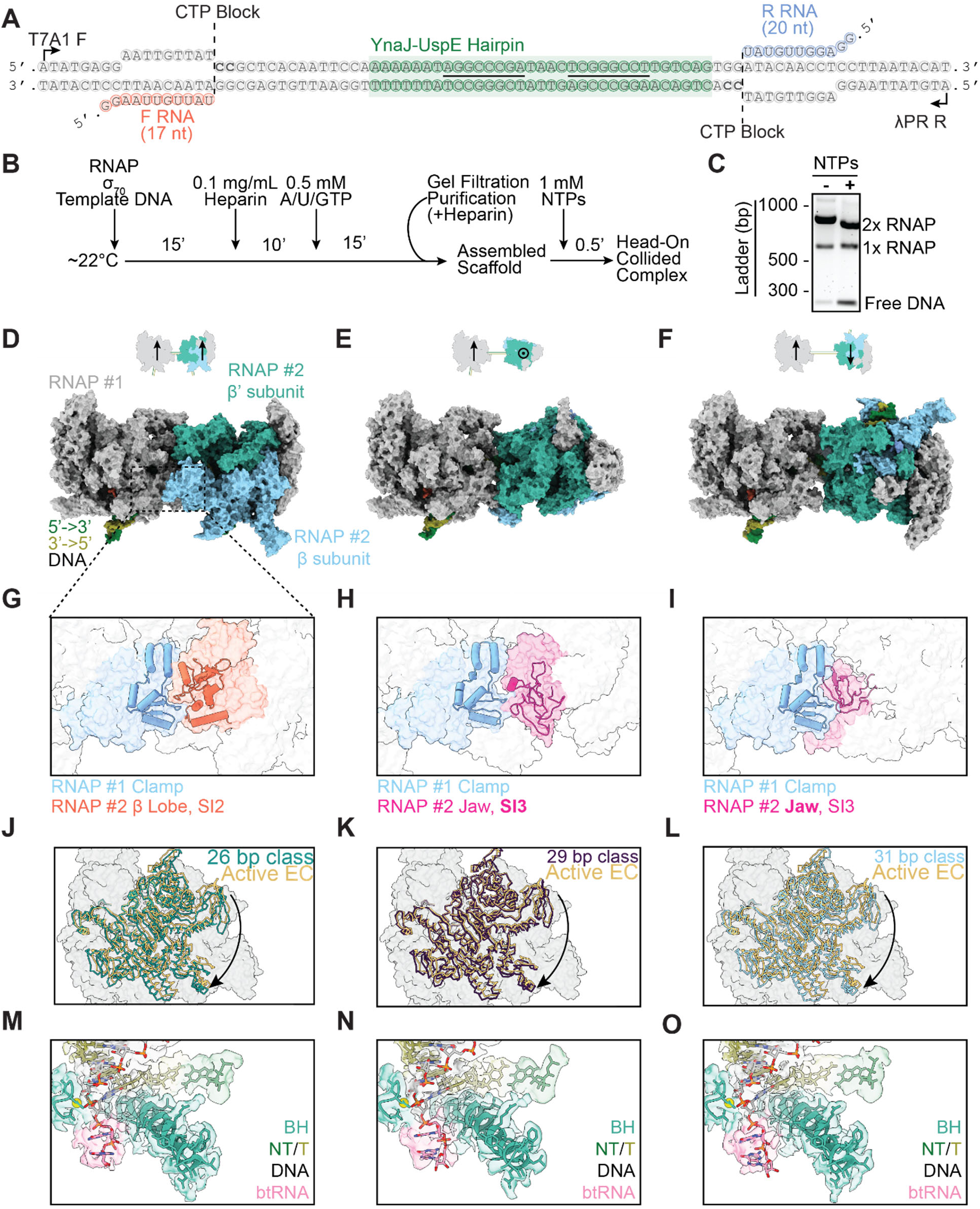
Reconstituting head-on RNAP-RNAP collisions. **[A]** DNA template featuring two promoters, designed to stall RNAP at cytosine sites. F RNA, forward RNA; R RNA, reverse RNA. Underlined segments of the YnaJ-UspE hairpin indicate complementary regions. **[B]** Experimental scheme for head-on RNAP-RNAP collided complex assembly and purification. **[C]** Native gel shift assay for evaluating collided complex formation. **[D-O]** Columns correspond to 26-bp (left), 29-bp (middle), and 31-bp (right) DNA inter-RNAP distance complexes. **[D-F]** Top: Cartoons indicating the relative orientation (arrows) and position of polymerases. Bottom: Surface representations of the corresponding atomic models. **[G-I]** Zoom-in views of the inter-RNAP interface for each class. **[J-L]** Views of the swivel module for each class (from symmetry-expanded reconstructions) superimposed on an active EC structure after alignment. **[M-O]** Zoom-in views of the backtracked RNA (btRNA). Cryo-EM maps and atomic models are displayed.

After biochemically confirming the formation of a head-on RNAP-RNAP collided complex (**Fig. 4C**), we employed this DNA template (once again supplemented with 1.5 mM Fos-Choline-8 detergent) for cryo-EM sample preparation. Initial 2D class averages displayed features consistent with two RNAPs in close proximity (**Fig. S6**). Unlike the RNAP-EcoRI* collided complex, 3D classification of the RNAP-RNAP collided complex revealed substantial heterogeneity. Notably, the distance between the two RNAP molecules varied, with the best resolved classes featuring 26, 29, and 31 bp of DNA between the active sites of the polymerases (**Fig. 4D-F**). Because of the ∼10.5-bp helical pitch of B-form DNA, these three configurations all feature approximate C2 symmetry, a plausible explanation for why both RNAPs could be effectively aligned and averaged when separated by these particular distances. We exploited this C2 pseudosymmetry to perform symmetry expansion and analyzed all RNAP protomers from a given inter-RNAP distance class within a single frame of reference, facilitating high-resolution refinement (26 bp, 2.65 Å; 29 bp, 2.87 Å; 31 bp, 2.62 Å global resolution) (**Fig. S7**).

We could not interpret nucleic acid sequences in the density maps (which were modeled as homopolymers), presumably due to heterogeneity in where collisions occur along the DNA template. An additional potential source of heterogeneity in our preparation is the presence of σ^70^ to promote initiation, but the 31-bp class was the only one that featured weak density (only visible at low map contour levels) for the σ^70^ domain 2 bound to RNAP. Therefore, the variable inter-RNAP spacing we recovered through classification likely reflects inherent heterogeneity in RNAP-RNAP collisions. We once again employed 3DFlex to further improve the resolution of these classes and analyzed their conformational landscapes (**Fig. S7**).

The RNAP-RNAP interface varies considerably among the three classes due to the differential translational and rotational relative positions of the RNAP pair on the helical DNA template. Nevertheless, in all three classes, the clamp domain of one RNAP was engaged with different domains of the other collided RNAP, notably the β lobe/SI2 domain (26 bp class), SI3 domain (29 bp class), and jaw domain (31 bp class) (**Fig. 4G-I**). We reasoned this could influence RNAP swiveling, as the clamp domain is a component of the swivel module. Indeed, all of the RNAPs in head-on collided complexes adopted a swiveled conformation (26 bp, 3.9°; 29 bp, 2.3°; 31 bp, 4.0° relative to PDB: 6RH3) (**Fig. 4J-L**). Additionally, despite sequence ambiguity, all the RNAPs showed clear density for backtracked RNA in their secondary channel regardless of inter-RNAP distance (**Fig. 4M-O**). Thus, despite substantial differences at the collision interface, RNAP-RNAP collisions populate a similar conformational landscape as the RNAP-EcoRI* collision, suggesting a fundamental RNAP response to roadblocks.

### RNA hairpin impacts collision positioning and stability

To evaluate the bulk transcriptional output of the paired RNAPs in this system, which also serves as a proxy for their positioning along the DNA, we performed in vitro SEnd-seq (simultaneous 5’ and 3’ end RNA sequencing) to systematically profile end-to-end RNA products^17,46^. End-biotinylated DNA templates were used in these experiments to pull down collided complexes, enriching for RNAs that remained bound to RNAP. We also constructed a template without the hairpin-encoding sequence for comparison. The sequenced RNAs displayed homogeneous 5’ ends (corresponding to initiation from either promoter) and more heterogeneous 3’ ends (**Fig. 5A, B, S8**), as anticipated given the stochastic nature of RNAP restart^47^, which will impact the occurrence and location of collisions. Samples including the hairpin-encoding sequence in the template featured a pronounced drop in the RNA length profile at the predicted hairpin sequence boundaries (**Fig. 5A**), resulting in significantly shorter transcripts for both the forward and reverse directions compared to the no-hairpin template (**Fig. 5C, S8**). These results indicate that the hairpin fosters consistent positioning of RNAP-RNAP collisions, mitigating heterogeneity attributable to the stochastic mechanochemistry of these nanoscale machines.

**Figure 5.**
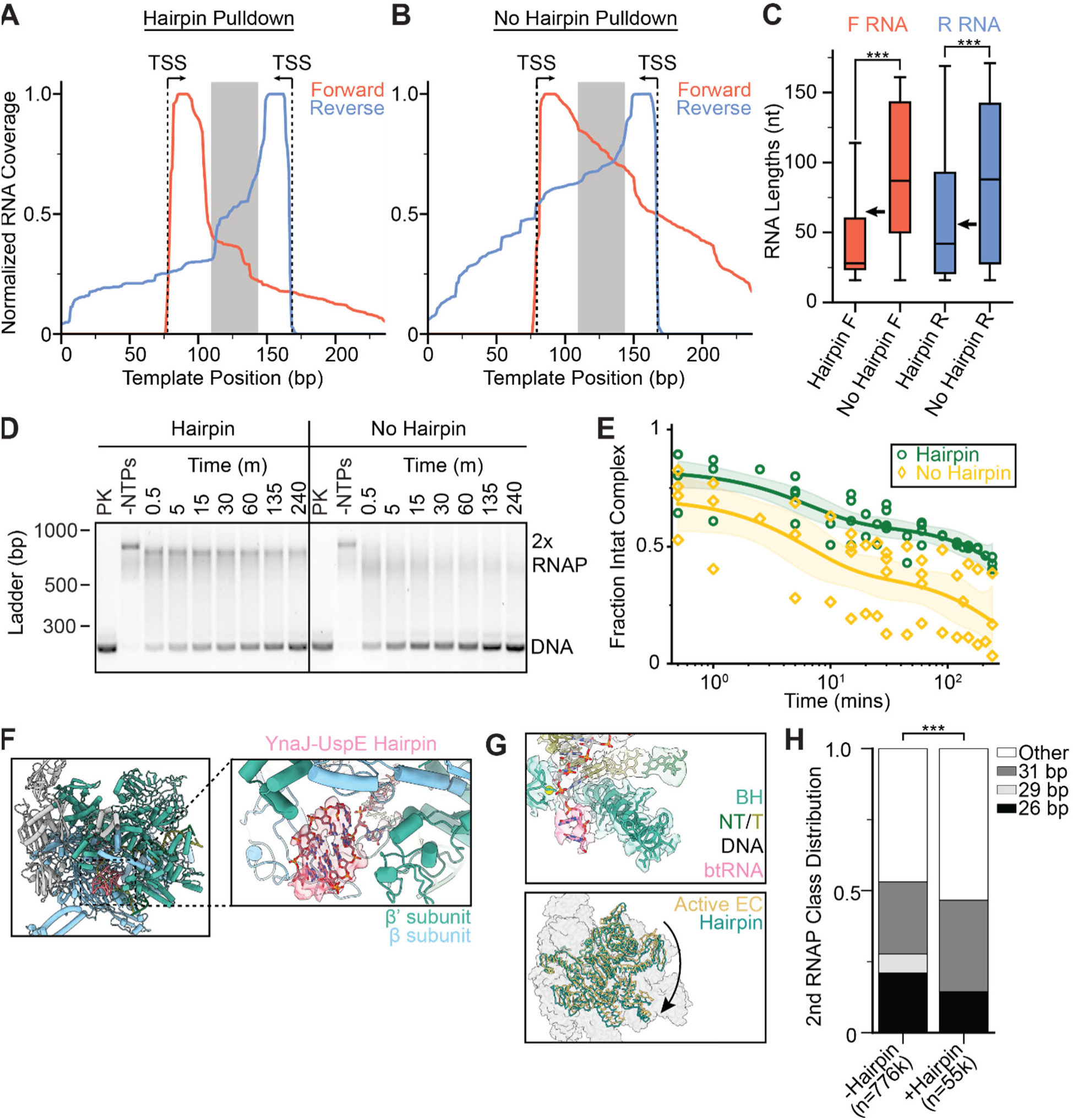
RNA hairpin impacts collision positioning and stability. **[A, B]** SEnd-seq profiles of forward and reverse RNAs in head-on RNAP-RNAP collided complexes on a template encoding the YnaJ-UspE hairpin (**A**) or a no-hairpin template (**B**). The grey box defines the hairpin sequence region or the corresponding scrambled sequence region. **[C]** Quantification of RNA lengths (mapped back to the originating promoter). N = 1789 (hairpin F), 8075 (no hairpin F), 608 (hairpin R), and 5945 (no hairpin R). Arrows indicate the predicted RNA lengths if transcription terminates right after the hairpin. Conditions were compared with a two-tailed, unpaired t test with Welch’s correction. ***p < 0.001. **[D]** Native agarose gels for evaluating collided complex disassembly. PK: Proteinase K treated. Timepoints indicate time elapsed after NTP addition, followed by EDTA quench. Left: hairpin-containing template DNA; right, no-hairpin template DNA. DNA was visualized with SYBR safe. **[E]** Quantification of the fraction of intact collided complex under indicated conditions. Nucleic acid intensity values were normalized for each of four experimental replicates to the -NTPs and +Proteinase K conditions. Shaded areas indicate 95% confidence interval of the fits. **[F]** View of the hairpin-containing head-on collided RNAP structure obtained through symmetry expansion. **[G]** Views of the hairpin-containing RNAP (cryo-EM map and atomic model) highlighting the backtracked RNA (top) and rotation of the swivel module relative to the active elongation complex. **[H]** Class distributions of no-hairpin and hairpin-containing collided RNAP complexes. “Other” is defined as particles that show well-defined density for one RNAP but the density downstream of the well-defined RNAP is too low of a resolution to confidently model an RNAP. Significance was calculated via a chi-squared test. ***p < 0.001.

We observed that the complex disassembles when incubated for longer time periods, which could occur when one RNAP is able to overcome the other and push it off the DNA template. Given the observation that all RNAPs in the collided complex were backtracked, we reasoned that the hairpin could serve as a barrier against long-range backtracking, thereby stabilizing the complex. To test this hypothesis, we performed a time course assay to measure the half-life of the collided complex by halting the transcription reaction with an EDTA quench at different timepoints post NTP addition. While the collided complex on the template with the hairpin remained stable for hours (Τ_1/2_ = 150 ± 40 min), the collided complex on the template without the hairpin disassembled within minutes (Τ_1/2_ = 6 ± 3 min) (**Fig. 5D, E**), confirming the hairpin’s contribution to complex stability.

We next sought to examine how the presence of a hairpin in the nascent transcript, which would form if one of the RNAPs has transcribed past the hairpin-encoding sequence before collision, influences the structure of the collided complex. No clear RNA hairpin density was observed in the exit RNA channel for any of the three RNAP-RNAP complex classes, indicative of flexibility or variable occupancy. We then reprocessed the data, re-extracting particles using a smaller box size encompassing one RNAP followed by focused classification at the RNA exit channel (**Fig. S9**). In a single class obtained using this approach (representing ∼7% of the data), we observed hairpin density (**Fig. 5F**). RNAP was still backtracked and swiveled (∼3.4°) in this class, suggesting that the presence of a well-ordered hairpin does not grossly alter RNAP’s conformational response to collision (**Fig. 5G**). To assess the hairpin’s impact on collision geometry, we recentered our particle picks to capture the other collided polymerase, allowing us to analyze the distribution of inter-RNAP spacings for particles contributing to the hairpin-containing class versus the distribution for the entire collided population. In the hairpin-containing particles, we found the class with the largest separation between RNAPs (31 bp) was mildly enriched (∼1.3-fold increase), while the classes with shorter inter-RNAP spacing were less populated, with a lower 26-bp class abundance (∼1.5-fold decrease) and a complete absence of the 29-bp class (**Fig. 5H**). These data are consistent with the notion that the hairpin reduces the heterogeneity of RNAP-RNAP collisions (**Fig. 5H**, p<0.001), potentially by restricting the dynamics of the hairpin-associated RNAP and maintaining its position on DNA.

### DNA deformation-RNAP swivel coupling depends on inter-RNAP distance

As a physical connection between RNAP and roadblock through the template DNA is a common element of transcriptional collisions with distinct composition and geometries, we reasoned this linkage could mediate characteristic RNAP remodeling. To test this hypothesis, we examined the coupling between DNA deformation and RNAP swiveling across RNAP-RNAP collision configurations. Swiveling was present in all three head-on collision orientations. However, interestingly, the degree of RNAP swiveling varied between the different classes in a manner that was non-monotonic with inter-RNAP distance, as the 29-bp spaced class featuring a substantially lower swivel angle in the consensus class (∼2.3°) than the 26-bp (∼3.9°) and 31-bp (∼4.0°) classes (**Fig. 4J-L**). We speculate this variation is linked to the differential interfaces between the RNAPs, which likely impact the local mechanical environment. We also found that the degree of DNA deformation downstream of the collided RNAP varied among classes (**Fig. 6A**). Finally, we examined whether the RNAP swivel angle and the DNA deformation remain correlated across different collision scenarios. When compared alongside previous RNAP structures representative of active EC, backtracked, and swiveled complexes (PDB: 6RH3, 6ASX, 6RI9, 6RIP)^35,39^, changes in the DNA PC1-axis did not correlate with changes in swiveling while changes in the DNA PC2-axis displayed a strong correlation, with the displacement increasing linearly with the magnitude of the RNAP swivel angle (**Fig. 6B, C**). Additionally, the extent of PC2 displacement within individual 3DFlex variability trajectories also correlates with the magnitude of swivel angle change (**Fig. 6D, S10, Movie S1, 3**)^30^. These analyses highlight that while RNAP’s conformational landscape varies across collisions, likely due to variations in the details of the RNAP-roadblock interface restricting the accessible DNA bending landscape, DNA deformation serves as a common mechanism to transduce collisions into RNAP swiveling and inactivation.

**Figure 6.**
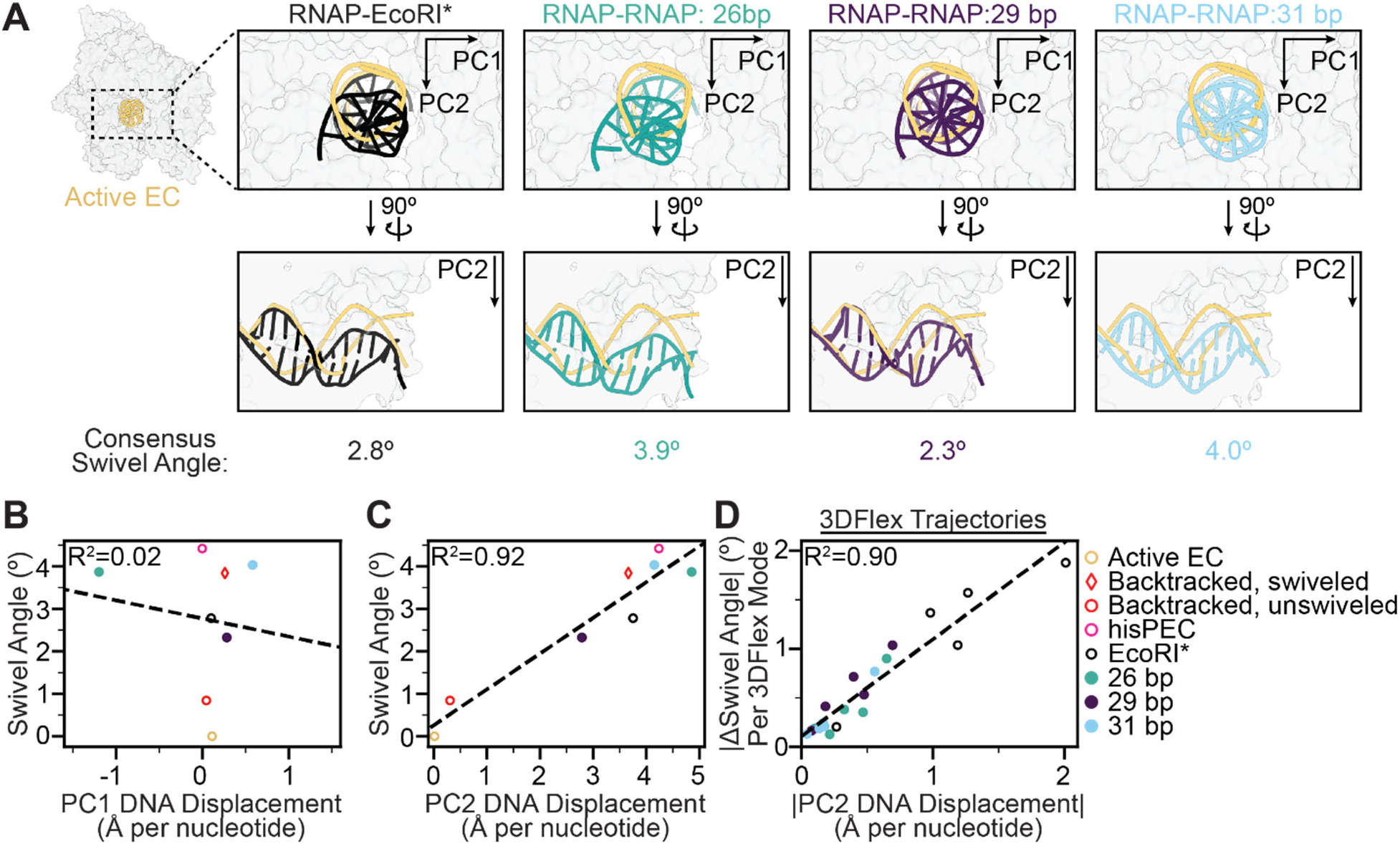
Inter-RNAP distance modulates deformation-swivel coupling. **[A]** Downstream DNA deformation of each collision scenario compared against active EC (PDB 6RH3). **[B-C]** Plots of swiveling angle versus DNA displacement along **[B]** PC1 and **[C]** PC2 of consensus collision structures (this study) and previously reported structures of an active EC (PDB 6RH3), backtracked ECs (unswiveled, PDB 6RI9; swiveled, PDB 6RIP), and his pause EC (PDB 6ASX). Linear fits are displayed (dashed lines). **[D]** Plot of the magnitudes of swiveling and PC2 displacement measured in individual 3DFlex series trajectories for the four collided structures (EcoRI*, 26 bp, 29 bp, 31 bp – 5 series each). A linear fit is displayed (dashed line).

## Discussion

In this study, we visualized the structural dynamics of RNAP upon collisions with both a passive (EcoRI*) and an active (another RNAP) roadblock under non-equilibrium, actively transcribing conditions. Despite variability in the composition and geometry of these collisions, our results reveal a characteristic RNAP response consisting of backtracking and swiveling to enter a transcriptionally inactive state (**Fig. 7A**). Our data highlight how shared mechanical elements across collisions, including roadblock steric hindrance and DNA deformation, can coordinate this common RNAP response irrespective of the detailed stereochemistry of a collision.

**Figure 7.**
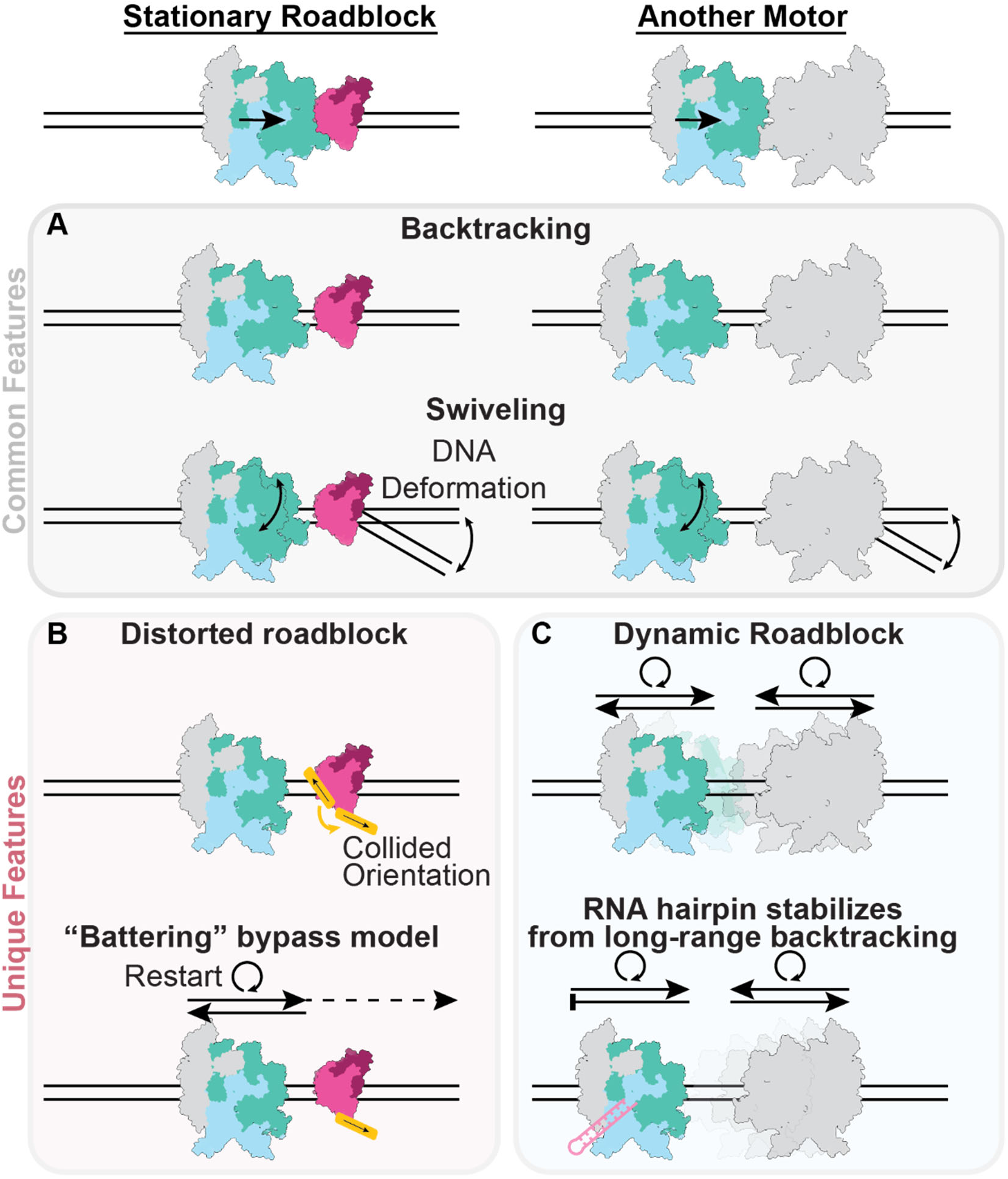
Structural mechanisms of transcriptional collisions. **[A]** Common features of all collisions in this study. Regardless of the roadblock identity, RNAP backtracks away from the collision site and swivels into a transcriptionally inactive state, mediated by coupling between template DNA distortion and RNAP swiveling. **[B, C]** Unique features of different collisions. **[B]** RNAP collision with a model stationary roadblock (EcoRI*) features roadblock remodeling, a plausible mechanism for how ‘battering’ (repetitive transcriptional restart and collision) can facilitate roadblock bypass. **[C]** Dynamic RNAP-RNAP collisions feature substantial heterogeneity due to the stochastic transcriptional and backtracking activity of both polymerases. These dynamics can be tempered by an RNA hairpin that constrains the collision positions.

As we did not directly visualize RNAP at its site of closest contact with roadblocks (indicated from transcript lengths), we cannot comment on how backtracking is triggered. We nevertheless speculate that roadblock steric hindrance ultimately limits forward progress of RNAP, triggering backtracking to a metastable position where clashes are moderated. At this position, RNAP dynamics can be modulated by coupling between DNA template deformation and RNAP swiveling, facilitating the maintenance of an inactivated state. Consistent with a previously proposed battering model^15,24^, we find that factors which reduce RNAP swiveling and enhance transcription restart after backtracking enhance roadblock bypass (**Fig. 3B-C, 7B**). Our structures also rationalize the previous observation that applying small opposing forces (∼0.2 pN) counterintuitively increases the rate of EcoRI* bypass by RNAP^24^, as this likely promotes RNAP un-swiveling by altering the downstream DNA deformation profile to more efficiently restart elongation after backtracked RNA cleavage, leading to eventual bypass. We furthermore uncovered an additional layer of the battering mechanism, wherein RNAP can physically deform roadblocks to eventually displace them (**Fig. 3D-F, 7B**). This suggests mechanical destabilization of deleterious roadblocks plays an instrumental role in their clearance from the genome.

Our observation that RNAPs in head-on collisions are also backtracked and swiveled, but feature variable inter-RNAP distances, is compatible with a scenario featuring intermittent, ping-ponging restart dynamics between the two RNAPs. We propose that when one RNAP stochastically restarts, it backtracks upon colliding with the other paused RNAP. This clears the path for either RNAP to once again restart, generating cycles of randomly alternating transcription and backtracking (**Fig. 7C**). This model can explain our previous observation that both RNAPs can transcribe through bidirectional convergent terminators like YnaJ-UspE before termination^17^, which is otherwise incompatible with a static collision. Our analysis also provides a mechanistic explanation for the positioning effect of the RNA hairpin when bound to RNAP, as we find its presence is correlated with increased complex stability (**Fig. 5D-E**) and reduced heterogeneity in inter-RNAP spacing (**Fig. 5H**). This suggests that the hairpin likely suppresses the motions of the engaged RNAP, thereby stabilizing it in a particular region along the DNA template and attenuating the variability in transcript length. We did still observe substantial variability in transcripts even in the presence of the hairpin in our minimal reconstitution system (**Fig. 5A-C**), suggesting additional cellular factors may further enhance the robustness of this mechanism, an important topic for future studies.

This work establishes the feasibility of scrutinizing genomic collisions using cryo-EM, providing a path towards visualizing physiological scenarios of increasing complexity. An important consideration for future studies is the topology of the template DNA, which may be addressable by employing closed templates with varying degrees of supercoiling^48,49^. Additionally, collisions featuring additional machines, such as co-directionally transcribing RNAP and translating ribosomes (both of which can promote roadblock bypass^3,10,15^) or head-on/co-directional DNA replication forks (replication-transcription conflict as a primary source of genome instability^4–6^) are likely accessible by similar approaches. Such studies will be critical for a mechanistic understanding of how mechanical conflicts are managed across the crowded genome and functionally harnessed to enhance the robustness of gene expression.

## Methods

### Protein purification

#### E. coli RNA Polymerase core

RNAP was expressed and purified as described previously^50^, with an additional step of 3C cleavage of the β’-3C-His10 tag and a reverse Nickel-IMAC (immobilized metal affinity chromatography) column to isolate RNAP core without the His10 tag.

#### Maltose Binding Protein (MBP)-EcoRI* (E112Q) and additional mutants

EcoRI* E112Q (a gift from Dr. Ruben Gonzalez’s lab) was cloned into a pET21 vector with an N-terminal His10-MBP-3C sequence using Gibson assembly. The plasmid was transformed into competent Eco BL21(DE3) by heat shock. The cells were grown in the presence of 50 µg/mL kanamycin to an optical density (OD) at 600 nm of ∼0.5-0.6 in a 37 °C shaker. Expression was induced with addition of 0.5 mM Isopropyl β-D-1-thiogalactopyranoside (IPTG) at 16 °C overnight. Cells were harvested by centrifugation, resuspended in lysis buffer [20 mM potassium phosphate, 300 mM sodium chloride (NaCl), 1 mM dithiothreitol (DTT), 0.1 mM ethylenediaminetetraacetic acid (EDTA), 5% glycerol (v/v), 0.15% Triton X-100, 1x protease inhibitor cocktail (Sigma-Aldrich), pH 7.5 at room temperature], and flash frozen in liquid nitrogen prior to storage at -80 °C.

Cell pellets were thawed, then lysed with an Avestin Emulsiflex C5 homogenizer. The lysate was then clarified (2x centrifugation steps, 30 minutes each, 48,000 x *g*) and applied to a 5 mL MBPTrap HP column (Cytiva). The column was washed with MBP Buffer A [20 mM potassium phosphate, 300 mM NaCl, 1 mM DTT, 0.1 mM EDTA, 5% glycerol (v/v), 0.15% Triton X-100, pH 7.5 at room temperature] and eluted with MBP Buffer A supplemented with 10 mM maltose. Peak fraction were pooled, diluted to in Heparin Buffer A [20 mM potassium phosphate, 1 mM DTT, 0.1 mM EDTA, 5% glycerol (v/v), 0.15% Triton X-100, pH 7.5 at room temperature] to 50 mM NaCl final concentration and loaded onto a 10 mL Heparin HP (Cytvia) column equilibrated in 10% Heparin Buffer A/90% Heparin Buffer B [20 mM potassium phosphate, 1 M NaCl,1 mM DTT, 0.1 mM EDTA, 5% glycerol (v/v), 0.15% Triton X-100, pH 7.5 at room temperature]. The bound protein was eluted with a gradient of 10-100% Heparin Buffer B over 10 CVs. Peak fractions of protein without nucleic acid contamination were eluted in a peak centered at ∼45 mS/cm conductivity and concentrated with a 30 kDa molecular weight cut-off (MWCO) concentrator (Amicon). Concentrated protein was clarified (5 minutes, 21,000 x *g*) before application onto a HiLoad 16/600 Superdex (Cytiva) gel filtration column equilibrated with gel filtration buffer [20 mM potassium phosphate, 300 mM NaCl, 1 mM tris(2-carboxyethyl)phosphine (TCEP), 0.1 mM EDTA, 5% glycerol (v/v), 0.15% Triton X-100, pH 7.5 at room temperature]. Peak fractions corresponding to the EcoRI* dimer were pooled, concentrated in a 30 kDa MWCO concentrator (Amicon), diluted with gel filtration buffer supplemented with 50% glycerol to a final concentration of 20% glycerol, flash frozen in liquid nitrogen, and stored at −80°C until use. Protein concentration was estimated using a Bradford assay^51^ (Bio-Rad Protein Assay Dye Reagent Concentrate) with bovine serum albumin standards.

For single point mutants (I21A, L108A, V166A, F252A, I258A), the same protocol was used with the following substitutions: 1 mM DTT was replaced in all buffers with 1 mM TCEP, and a Superdex 200 Increase (Cytiva) column was used for gel filtration instead of a HiLoad 16/600 Superdex (Cytiva) column.

#### GreA, GreB, NusA, NusG, σ^70^

GreA, GreB, NusA, NusG, and *σ*^70^ were expressed, purified was performed as described previously^16,50,52,53^.

Endonucleolytic cleavage activity of GreA/B was verified as described previously^16^. In brief, a backtracked RNAP EC scaffold (nucleotide sequences in **Table S1**) was formed with a short (2 nt RNA backtracked) and long backtracked EC (6 nt RNA backtracked). Oligonucleotides were high-performance liquid chromatography (HPLC) purified (IDT). Activity assays were performed at 37 °C for 10 minutes by adding GreB or GreA at a final concentration of 600 nM in the absence or presence of NTPs. The reactions were stopped by addition of an equal volume of 2x RNA loading buffer and incubation at 90 °C for 2 minutes. Products were separated on a 25% Urea-PAGE gel, followed by fluorescence visualization on a Typhoon FLA 7000 scanner. Full-length RNA products indicative of endonucleolytic cleavage and transcriptional restart were enhanced in the presence of GreA (short backtracked EC) and GreB (long backtracked EC) as expected^33^.

As reported previously, NusA and NusG were verified to be active by bulk in vitro transcription assays in transcription buffer (25 mM Tris-hydrochloride pH 7.5, 150 mM potassium chloride, 10 mM magnesium chloride, 1mM DTT). In brief, Rho and NusG in combination were shown to enhance Rho mediated termination efficiency on the Rho-dependent λtR1terminator^53^. NusA reduced transcriptional readthrough across the ynaJ-uspE bidirectional terminator^54^. σ^70^ was verified to be active in promoting T7A1 and LambdaPR dependent promoter complex formation and transcription in in vitro transcription assays in a transcription buffer (25 mM Tris-HCl pH 7.5, 150 mM KCl, 10 mM MgCl_2_, 1mM DTT)^55^.

## RNAP-EcoRI* Scaffold Complex Assembly

DNA oligonucleotide sequences (Polyacrylamide gel electrophoresis (PAGE)-purified from IDT) and a 5’ Cy3-labeled RNA oligonucleotide (Dharmacon) were designed using a previously used scaffold as a starting point (sequences of template, non-template, and RNA oligos in **Table S1**)^56^. Template DNA and RNA were annealed in a 1:1 ratio in 1x IDT annealing buffer (100 mM potassium acetate (KOAc); 30 mM HEPES, pH 7.5) for 5 minutes at 95⁰ C, 5 minutes at 75⁰ C, 5 minutes at 45⁰ C, and 40 cycles of 1⁰ C steps (1 minute hold each step, final temperature 4⁰ C) in a thermocycler. RNAP core desalted into scaffolding buffer (20 mM Tris pH 7.5 at RT, 150 mM potassium glutamate (KGlu), 5 mM magnesium acetate (Mg(OAc)_2_), 2.5 mM DTT) was then incorporated and incubated for 15 minutes at RT. 1.5-fold excess of non-template DNA was then added to the complex mixture and incubated for 10 minutes at RT. 5-fold excess of EcoRI* was then added to the mixture (this step was excluded for the -EcoRI* controls) and incubated for 10 minutes at RT before addition of 0.1 mg/ml final concentration of heparin sodium salt to remove non-specifically bound RNAP and EcoRI* proteins. The fully assembled complex was purified away from incompletely assembled complexes via gel filtration chromatography using a Superdex 200 Increase column equilibrated with low salt buffer (20 mM Tris-acetate (Tris-OAc) pH 7.8 at RT, 50 mM KOAc, 5 mM Mg(OAc)_2_, 2.5 mM DTT) containing 0.1 mg/ml heparin (Sigma-Aldrich) to prevent re-binding of non-specifically loaded RNAP/EcoRI* to the DNA template during the purification. Complex assembly was monitored via native gel electrophoresis on a 5% polyacrylamide 0.5X TBE gel.

## RNAP Head-On Complex Assembly

DNA constructs featuring either two RNAP promoters (T7A1 and λPR) or only one promoter were generated by PCR, purified via gel filtration chromatography on a Superdex 200 Increase column equilibrated with DNA gel filtration buffer (20 mM Tris pH 8.0 at RT, 200 mM NaCl, 0.1 mM EDTA), and concentrated into Tris Low EDTA buffer (10 mM Tris pH 8.0 at RT, 0.1 mM EDTA) for long term storage. PCR oligos and full DNA sequences of the templates are in **Table S1**.

RNAP holoenzyme was formed by incubating σ^70^ with RNAP core for 20 minutes at 37⁰ C in their purification buffers, using a 3:1 ratio of σ^70^:RNAP core. The complex was then desalted into transcription buffer (25 mM Tris-HCl pH 7.5 at RT, 150 mM KCl, 10 mM MgCl_2_, 1mM DTT), and its concentration measured using A280 and its calculated extinction coefficient. To form open promoter complexes, 4-fold excess of holoenzyme was added to the DNA template (final concentration ∼1-2 μM DNA) and incubated for 15 minutes at 37⁰ C. Non-specifically bound RNAPs were removed by incubation with 0.1 mg/mL heparin for 10 minutes at 37⁰ C. 0.5 mM ATP, UTP, and GTP were then added to the reaction mixture and incubated for 15 minutes at 37⁰ C to walk the promoter bound RNAPs out to the dual CTP nucleotide deprivation stall site. The fully assembled complex was purified away from incompletely assembled complexes via gel filtration chromatography using a Superdex 200 Increase (Cytiva) column equilibrated with low salt buffer (20 mM Tris-OAc pH 7.8 at RT, 50 mM KOAc, 5 mM Mg(OAc)_2_, 2.5 mM DTT) containing 0.1 mg/ml heparin to prevent re-binding of non-specifically loaded RNAP to the DNA template during the purification. Complex assembly was monitored via native gel electrophoresis on a 2% agarose 0.5X TBE gel.

### Cryo-EM

### Sample Preparation

After gel filtration, the complex was concentrated using Amicon 100 kDa concentrators and washed 7x (450 μL added per wash) into low salt buffer without heparin to dilute any remaining heparin still in solution to a negligible amount. The complex was then diluted with low salt transcription buffer supplemented with Fos-Choline-8 (Anatrace, final concentration 1.5 mM). 3 μL of sample was incubated with 0.5 μl of 7x NTP stock solution (3.5 mM NTPs in low salt transcription buffer plus 1.5 mM Fos-Choline 8) for one minute (EcoRI* complex) or 30 seconds (RNAP-RNAP) at room temperature before vitrification. Samples were prepared on glow-discharged (Gatan Solarus, O_2_/H_2_, 10 seconds) C-Flat Holey Carbon Film CF-1.2/1.3-4, 400 mesh gold grids at concentrations ranging from 10-14 μM for RNAP-EcoRI* and ∼4 μM for RNAP-RNAP. Grids were loaded into Leica EM GP apparatus, blotted for 4-5 seconds at 90% humidity at room temperature, and plunge frozen in liquid ethane.

For the RNAP-EcoRI* complex, the sample concentration was estimated by the A550 signal (Cy3 extinction coefficient of 150000 M^-1^ cm^-1^). For the RNAP-RNAP complex, sample concentration was estimated using A260 and A280 absorbance measurements using combined extinction coefficients from the individual components.

### Cryo-EM data acquisition and processing

For RNAP-EcoRI* complex, three different grids were imaged from two different sample preparations of the complex, frozen on different days. Datasets were collected on the same FEI Titan Krios G3i system operating at 300 kV and equipped with a Gatan K3 direct electron detector, BioQuantum energy filter, and a spherical aberration corrector using super-resolution mode. Frame sequences (“movies”) were recorded using the SerialEM software^57^ suite at a nominal magnification of ×81,000, corresponding to a calibrated pixel size of 0.86 Å at the specimen level (super-resolution pixel size of 0.43 Å per pixel)^58^. Each 1.4 s exposure was dose-fractionated across 35 frames, with a total electron dose of 47.3 e^−^ Å^−2^ (1.35 e^−^ Å^−2^ per frame), with defocus values ranging from −0.8 to −2.0 μm underfocus. Beam-image shift was used to collect 43,378 single exposures (4.6k, 17.8k, 23.8k exposures from the different grids) from nine holes in a 3-by-3 grid per each stage translation.

The RNAP-RNAP complex dataset was collected similarly, with the following alterations. Data were collected on a single grid on a FEI Titan Krios G2 operating at 300 kV and equipped with a Gatan K3 direct electron detector and BioQuantum energy filter. Movies were recorded at a nominal magnification of x105,000, corresponding to a calibrated pixel size of 0.847 Å at the specimen level (super-resolution pixel size of 0.4235 Å per pixel)^58^. The total electron dose was 48.79 e^−^ Å^−2^ (1.39 e^−^ Å^−2^ per frame). Beam-image shift was used to collect 20,080 single exposures.

### Cryo-EM data processing

All processing was performed using cryoSPARC^59^. Movies were aligned with Patch Motion correction (B-factor: 150), and contrast transfer function (CTF) parameters were estimated with Patch CTF using default settings. For each dataset, blob picker was initially used to pick particles from a 256-pixel box, followed by 2D and 3D classification on extracted particles to remove junk particles and RNAPs that did not restart. This initial particle stack was used to generate a preliminary reconstruction that was used to re-pick particles using a template picker approach to yield more particles. All of the picked particles were then re-extracted with a 400-pixel box and separated with multiple rounds of 3D heterogeneous refinement job with candidate 3D volumes as well as junk particle volumes (**Fig. S2, S6**). All models were then further refined via a non-uniform refinement job^60^, local CTF refinement, and global CTF refinement. Then low per-particle scale particles were removed, followed by per-particle reference-based motion correction and subsequent local refinement. 3DFlex^30^ was then performed (N=5 modes) to improve the density of high flexible regions in the particles and generate variability trajectories for DNA deformation-RNAP swiveling analysis.

For the RNAP-RNAP dataset, the following additional steps were performed. During test dataset processing a 400-pixel box size was used and binned by 4 to enhance processing speed during initial processing steps. Particle box size was increased to 700 pixels (no binning) to accommodate the additional RNAP. After heterogeneous refinement, an additional round of 3D classification was performed using low-resolution volumes as bait volumes (generated by manual docking to mimic RNAPs with 26-36 bp of separation followed by molmap (resolution 20) volume generation from atomic models) using ChimeraX to see if any additional states could be extracted. This resulted in the detection of the 29 bp inter-RNAP class. After non-uniform refinement, models were re-oriented along their C2 symmetry axis and symmetry expanded before local refinement and further polishing with a one RNAP sized mask.

To isolate particles that had clear hairpin density in the exit channel, a new set of particles were extracted from the same micrographs (to remove any potential bias from the C2 symmetric particles that were able to be resolved to high resolution) and subjected to a similar processing pipeline to a resolve a single RNAP class (box size 400 pixels) using a focus mask corresponding to a single RNAP (**Fig. S9**). This class contained downstream DNA and low-resolution signal further downstream that could correspond to collided RNAP density being averaged between different states (since the second RNAP was not included in the focus mask). Taken together, this indicates that these single RNAPs are still in a post-collision state instead of the alternate hypothesis such as run-off RNAPs or non-bound RNAPs. These particles were subjected to polishing and refinement prior to an additional round of focused classification performed with a mask around the RNAP exit channel that showed one class with clear RNA hairpin density. Particles contributing to the hairpin-containing class were subjected to final polishing and refinement. Additionally, to compare the collided states of the hairpin-containing RNAPs, hairpin-containing particles were re-extracted with a larger box size (700 pixels) and subjected to 3D classification with a focus mask excluding the high-resolution hairpin-containing RNAP (N=8 classes). The relative distribution of classes representing either the 26, 29, 31 bp inter-RNAP distances, or other classes consisting of only a single well-refined RNAP was then determined via manual inspection of the density maps and comparison to the 26, 29, and 31 bp maps constructed prior.

### Model building and refinement

The RNAP-EcoRI* initial model was derived from PDB 6RIP^35^. The model was manually fit into the cryo-EM density maps using ChimeraX^37^, modified in Coot/PyMOL/Isolde^61–63^, and followed by real-space refinement using PHENIX^64^. The nucleic acid sequence was fixed in the same reference frame as the EcoRI* cognate binding sequence, which matched the cryo-EM map along the rest of the nucleic acid sequence. For real-space refinement, rigid body refinement was followed by all-atom refinement with Ramachandran and secondary structure restraints. For the RNAP-RNAP complexes, the same refinement methodology was applied with the following modifications. The RNAP coordinates from the RNAP-EcoRI* complex were used as the starting model for the RNAP-RNAP complexes. Additionally, since we were unable to determine the nucleic acid sequence by visual inspection in any of the RNAP-RNAP states (likely due to heterogeneity in the positioning of collision sites along the template), poly-G and poly-C sequences were used during modeling building.

### Computational modeling of collision interface

The nucleic acids of the RNAP-EcoRI* collision were converted to homopolymer models to make modeled transcription steps (stepwise rotation and translation along the DNA axis) easier. The RNAP coordinates were transformed in three-dimensional space by stepwise alignment of the downstream DNA nucleotides to the same sequences plus one step further downstream. SES overlap volume was calculated by summation of the SES volumes of the individual RNAP and EcoRI* proteins and subtracting the SES volumes for the combined RNAP and EcoRI. In the case where the RNAP and EcoRI* volumes are overlapping, their combined surface volumes will be less than the sum of the individual protein volumes yielding a positive SES overlap volume. SES overlaps were re-scaled such that the 39 nt position equaled zero. Alignment and calculation of SES volumes were performed in ChimeraX^37^.

## 3DFlex model fitting individual frames

For each frame of the 3DFlex variability trajectories, docking of individual rigid bodies (as defined in **Table S3**) from the consensus model was performed in ChimeraX^37^ followed by subsequent unsupervised all-atom refinement in Isolde (5 minutes, 20K, nucleotides restrained by base-pair hydrogen bonding)^63^.

### Swivel domain motion calculation

The core modules of the different RNAP structures were superimposed in PyMOL by aligning the following the α-carbons: all modeled residues from the α subunit, residues 3:27, 142:152, 445:455, 520:713, 786:828 from the β subunit, residues 343:368, 412:524, 530:552, 569:701, 720:786 from the β’ subunit, and all modeled residues from the ω subunit.

RNAP EC (PDB: 6RH3)^35^ was used as the reference structure (swivel angle = 0°) for swivel angle measurements. To determine the swivel angle for a given RNAP structure, residues comprising the swivel module from the RNAP structure and the reference were selected (β-subunit AA 1241:1341; β’-subunit AA 1:342, 369:420, 787:930, 1135:1375). The rotation axis and rotation angle transforming one swivel module into the other were then computed using the PyMOL script draw_rotation_axis.py (https://pymolwiki.org/index.php/RotationAxis). For DNA deformation-RNAP swiveling coupling analysis, the same method was applied. For symmetry expanded particles in the head-on collision state, the swivel angle of only the RNAP contained in the refinement mask was analyzed.

### DNA displacement calculation

Generation of a frame of reference was necessary to orient downstream DNA displacement on different axes. To accomplish this, the center of masses of twelve downstream nucleotides of the active RNAP EC (PDB: 6rh3)^35^ were extracted using PyMOL^62^. PCA was performed on their 3D coordinates to generate the PC1 and PC2 orthoplanes along the long axis of the downstream DNA. Since this definition of PC1 and PC2 is due to inherent variation of the center of mass of the nucleotides downstream of the active site in the reference frame RNAP EC (PDB: 6rh3)^35^, the plane definition is arbitrary, meaning that the definition of which axis is most responsible for variance (PC1) is descriptive of the RNAP reference structure variability, not descriptive of the variance between the different structures being compared in this study. For each query structure (consensus structures and 3DFlex frames), the structural cores of the RNAPs were aligned to the structural core of the active RNAP EC (PDB: 6rh3)^35^, and displacement of the center of mass of each nucleotide was then projected onto PC1 and PC2, summed across the sequence of twelve nucleotides, and averaged.

## Biochemical Assays

### RNAP-EcoRI* Collided Complex

#### Backtracking assays

5’ Cy3-labeled RNA oligonucleotides corresponding to zero to four nucleotides of backtracking (based on the RNAP starting position along the nucleic acid sequence) were synthesized (Dharmacon) and used as a ladder to verify the RNA length. For the backtracking assay, the scaffold was assembled as described previously except for omitting the EcoRI* binding step before gel filtration purification. Twenty-fold excess of EcoRI* (if applicable) and 1 μM Gre factors (if applicable) were added to the assembled complex and incubated at RT for 20 minutes before addition of 0.5 mM NTPs for 1 minute in a final volume of 15 μL. To process the RNA samples, 1.5 μL of Turbo DNase (ThermoScientific) was added and incubated for 15 minutes at 37⁰ C, 1.5 μL of proteinase K (New England Biolabs) and 50 mM EDTA for 15 minutes at RT, and finally an equal volume of 2x RNA loading dye (1x components - 47.5% formamide, 0.01% SDS, 0.01% bromophenol blue, 0.5 mM EDTA) was added to the samples. Samples were boiled at 95⁰ C for 5 minutes and separated on an 18% TBE-Urea PAGE gel and visualized using Cy3-fluorescence on a Typhoon gel imager. Three biological replicates of this assay were performed on different days.

#### EcoRI* bypass assays

RNAP-EcoRI* containing scaffolds were purified as described previously. Samples were then diluted into heparin-free buffer to a final concentration of ∼200 nM as quantified by Cy3 RNA fluorescence. Samples were then diluted two-fold into a mixture of low salt transcription buffer supplemented with varying KOAc concentrations and 0.5 mM NTPs. These reactions were incubated at 37⁰ C for 30 minutes and then quenched by addition of 2X RNA loading dye (95% formamide, 0.02% SDS, 0.01% bromophenol blue, 1 mM EDTA) supplemented with KOAc to equalize the final salt concentration between conditions. For analysis, the intensity of the roadblock and scaffold band were background subtracted, measured in FIJI^65^, and normalized relative to each other (roadblock intensity divided by scaffold intensity, and additionally rescaled for each salt titration gel so that the 50 mM normalized roadblock intensity was set to E_RB_ = 1, and the background intensity was set to E_RB_ = 0). E_RB_ calculated at 300 mM KOAc (because that salt concentration had the largest dynamic range when all the factor addition experiments were compared) was rescaled so that the average value of the EcoRI* (“normalized E_RB_”) was equivalent to 1 for each sample condition (factor addition experiments in **Fig. 3C** and stability mutants experiments in **Fig. 3F**) to adjust for observed batch effects when different NTP stocks were used.

### RNAP Head-On Collided Complex

#### Complex stability assay

RNAP-RNAP head-on complexes were assembled and purified for DNA templates with and without the YnaJ-UspE hairpin as previously described in low salt transcription buffer supplemented with 0.1 mg/ml heparin. Samples were concentrated using a 100 kDa MWCO concentrator (Amicon). Control samples to measure total DNA amount loaded per well were generated by treating samples with 1 µL proteinase K (New England Biolabs) per 10 µL reaction volume supplemented with 25 mM EDTA (final concentration) for over one hour at room temperature. 1 mM NTPs were added to initiate transcription restart, and aliquots of the reaction were quenched with addition of 25 mM EDTA (final concentration) at the specified time interval. Samples were run on a 2% 0.5X TBE agarose gel after being diluted with a 6x stock of 2M sucrose, and the resulting gel was stained with SYBR safe to visualize the DNA. Lane profiles were extracted using a 0.5x lane width window. Signal was summed in the 400-1000 bp range, then normalized between the -NTP control (corresponding to 100% RNAP-RNAP complexes) and the proteinase K control (corresponding to background signal) to calculate fraction intact complex. Four different experimental replicates were performed. Bi-exponentials were fitted through the data using the scipy python module^66^ to approximate the half-life of the collided complex, and 95% confidence intervals are displayed in the plot.

#### SEnd-seq

RNAP-RNAP head-on complexes were assembled and purified as previously described in low salt transcription buffer supplemented with 0.1 mg/ml heparin. For single promoter templates, the same general protocol was followed with a polymerase:DNA ratio of 2:1 instead of 4:1 as used for the double promoter templates. Samples were concentrated using a 100 kDa MWCO concentrator (Amicon). +PK labeled samples were treated with proteinase K (New England Biolabs) and 25 mM EDTA for over one hour at room temperature. 1 mM NTPs were added and incubated at RT for 1 minute before quenching with 25 mM EDTA. For complex pulldown, samples were washed and incubated with streptavidin M-280 Dynabeads (Thermo Fisher Scientific) before being pulled down via a magnetic separation rack (Permagen), then washed twice with 200 μL low salt buffer supplemented with 25 mM EDTA and 0.1 mg/mL heparin. The complexes were then treated with proteinase K for 1 hour at 37 °C to elute RNAs off the beads followed by overnight incubation at 4°C. The reaction was purified using 2.2x vol of Agencourt RNAClean XP beads (Beckman Coulter, A63987). After elution, nucleic acids were treated with 0.5 μl TURBO DNase (Life Technologies, AM2238) to remove template DNA. The remaining RNA was further purified via phenol-chloroform-isoamyl alcohol (Invitrogen, 15593031) extraction (25:24:1, v/v/v) and recovered by ethanol precipitation. The RNA pellet was resuspended, mixed with 1 µL of spike-in RNA, and subsequently ligated to a 3’ RNA adapter for SEnd-seq library preparation, as previously described^17,46^. Sequencing was performed on an Illumina MiSeq instrument in a paired-end mode (150 nt x2).

Reads were barcode sorted to the correct template and experimental condition. The separate 5’ and 3’ end reads of each RNA sequence were aligned to both the forward and reverse candidate RNA sequence using a local pairwise aligner in the Biopython module^67^ (match_score=2, mismatch_score=-1, open_gap_score=-5, extend_gap_score=-0.5). Alignment scores were calculated by the summation of the two best candidate positions. The higher scoring alignment was taken, as long as the distance between the 5’ and 3’ ends was greater than 15 nucleotides, as all the RNAs should be at least 17 nucleotides from the A/G/UTP only transcription initiation walkout before subsequent purification. Unmapped reads either had the wrong direction of 5’ and 3’ alignment or were too short. Furthermore, RNAs that did not initiate from within a 10-nucleotide range near the transcription start site were filtered out. These lost reads upon filtering could come from RNAP initiation from the end of the DNA template, RNA degradation of the 5’ end, or non-templated addition at the 3’ end^68^.

## Molecular graphics and data analysis

Molecular graphics were prepared with UCSF ChimeraX^37^. Plotting and statistical analysis were performed with Python using the matplotlib module^69^. Python codes were prepared with the assistance of ChatGPT 4.0.

## Statistical analysis

Conditions were compared via two-tailed, unpaired t-test including Welch’s correction. If applicable, a Benjamini-Hochberg correction for multiple hypothesis testing was added. S.D. stands for standard deviation and S.E. stands for standard error when used in the text. P-value asterisk reported as *p < 0.05, **p < 0.01, ***p < 0.001.

## Acknowledgements

We thank Johanna Sotiris, Honkit Ng, and Mark Ebrahim at the Rockefeller University Evelyn Gruss Lipper Cryo-EM Resource Center for their support in data collection. The expression plasmids for Rho, NusA, NusG, GreA and GreB were provided by Rachel Anne Mooney (Landick Laboratory, University of Wisconsin-Madison). NusA, NusG, GreA, and GreB proteins were purified by Rui Gong (Alushin Laboratory, Rockefeller University). J.W.W. acknowledges support from National Institutes of Health NRSA training grant T32GM066699. S.L. acknowledges support from National Institutes of Health grant R01GM149862, and G.M.A acknowledges support from grant R35GM161251. S.L. and G.M.A acknowledge support from the Stavros Niarchos Foundation Institute for Global Infectious Disease Research at the Rockefeller University and a joint award from the Alfred P. Sloan Foundation Matter-to-Life Program (G-2024-22709). G.M.A is a Biohub investigator.

## Author contributions

J.W.W., S.L., and G.M.A. designed research. J.W.W. purified proteins, collected cryo-EM datasets, and performed biochemical studies. A.U.M. purified proteins and helped with biochemical experimental design. A.U.M. and S.A.D. helped with cryo-EM processing and data interpretation. X.J. performed SEnd-Seq experiments. S.J.C. performed biochemical assays for the RNAP-EcoRI* complex. H.J.Y. performed biochemical assays for the RNAP-RNAP collided complex and collected the RNAP-RNAP collided cryo-EM dataset. J.W.W., S.L., and G.M.A. wrote the paper with input from all other authors.

## Data availability

Cryo-EM density maps and corresponding atomic models have been deposited in the PDB and EMDB with the following accession codes: RNAP-EcoRI* Collision (PDB: XXXX, EMDB: YYYYY); Head-on RNAP-RNAP Collision: 26 bp distance (PDB: XXXX, EMDB: YYYYY); Head-on RNAP-RNAP Collision: 29 bp distance (PDB: XXXX, EMDB: YYYYY); Head-on RNAP-RNAP Collision: 31 bp distance (PDB: XXXX, EMDB: YYYYY); Head-on RNAP-RNAP Collision: RNA hairpin (PDB: XXXX, EMDB: YYYYY). 3DFlex volumes and fit models for both consensus and variability trajectory movies have been deposited in zenodo (doi: 10.5281/zenodo.19317100).

## Competing interests

The authors declare no competing interests.

## Notes

https://zenodo.org/records/19317100

### Competing Interest Statement

The authors have declared no competing interest.

### Summary of Updates

Additional funding support included. Methods section added for computational modeling of collision interface.

