## Supplemental Figures and Tables for "Structural principles of transcriptional collisions"

### Supplementary Information

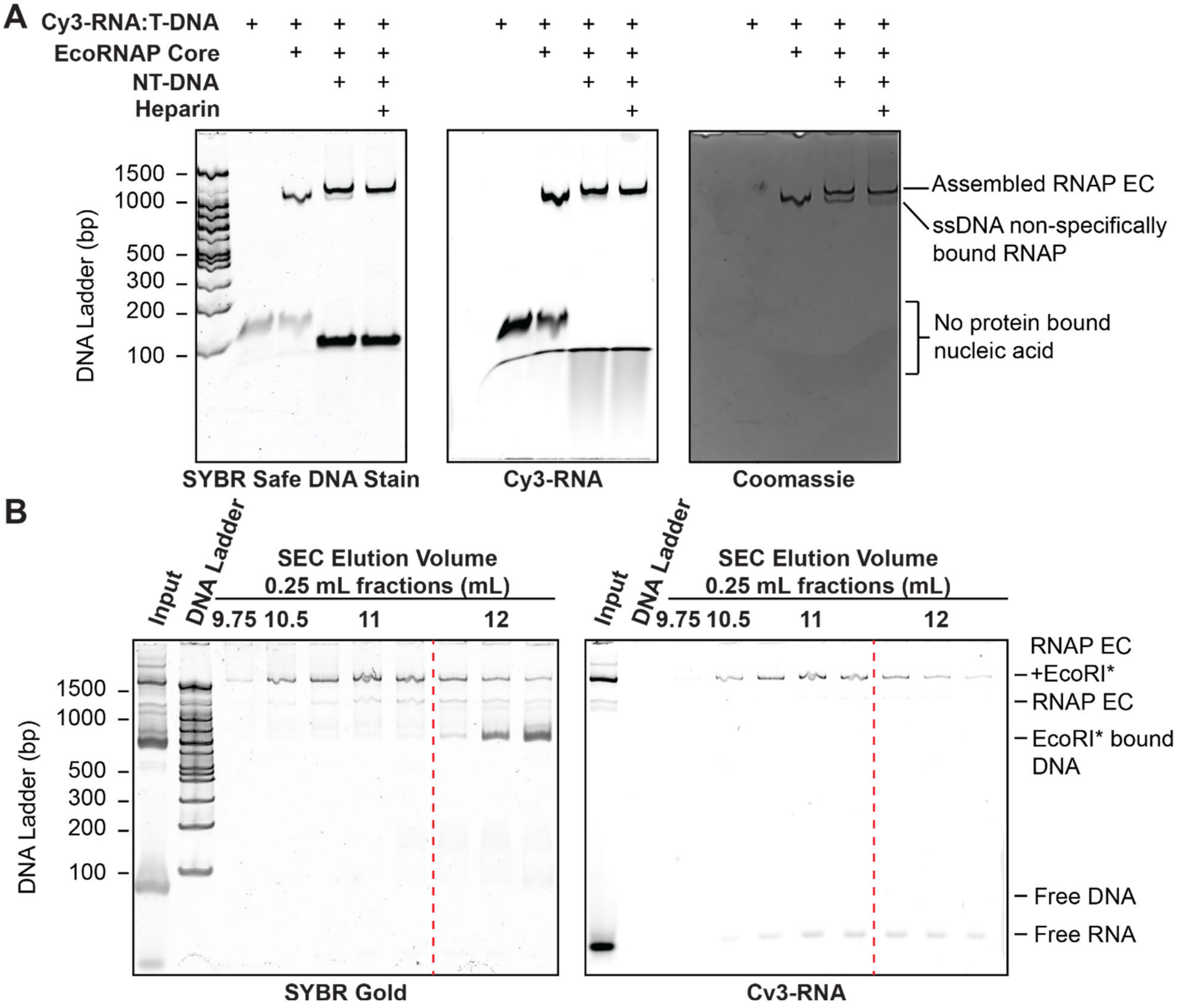


**Supplementary Figure 1. RNAP-EcoRI* collided complex assembly and purification.**

**[A]** Assessment of RNAP scaffold assembly prior to EcoRI* addition. Quality control staining for DNA (left), RNA (middle), and protein (right).

**[B]** Analysis of the RNAP-EcoRI*-DNA ternary complex after gel filtration, staining for DNA (left) and RNA (right). Samples were pooled from the left of the red dashed line and concentrated prior to freezing. 100 bp gold bio ladder was used for all experiments.

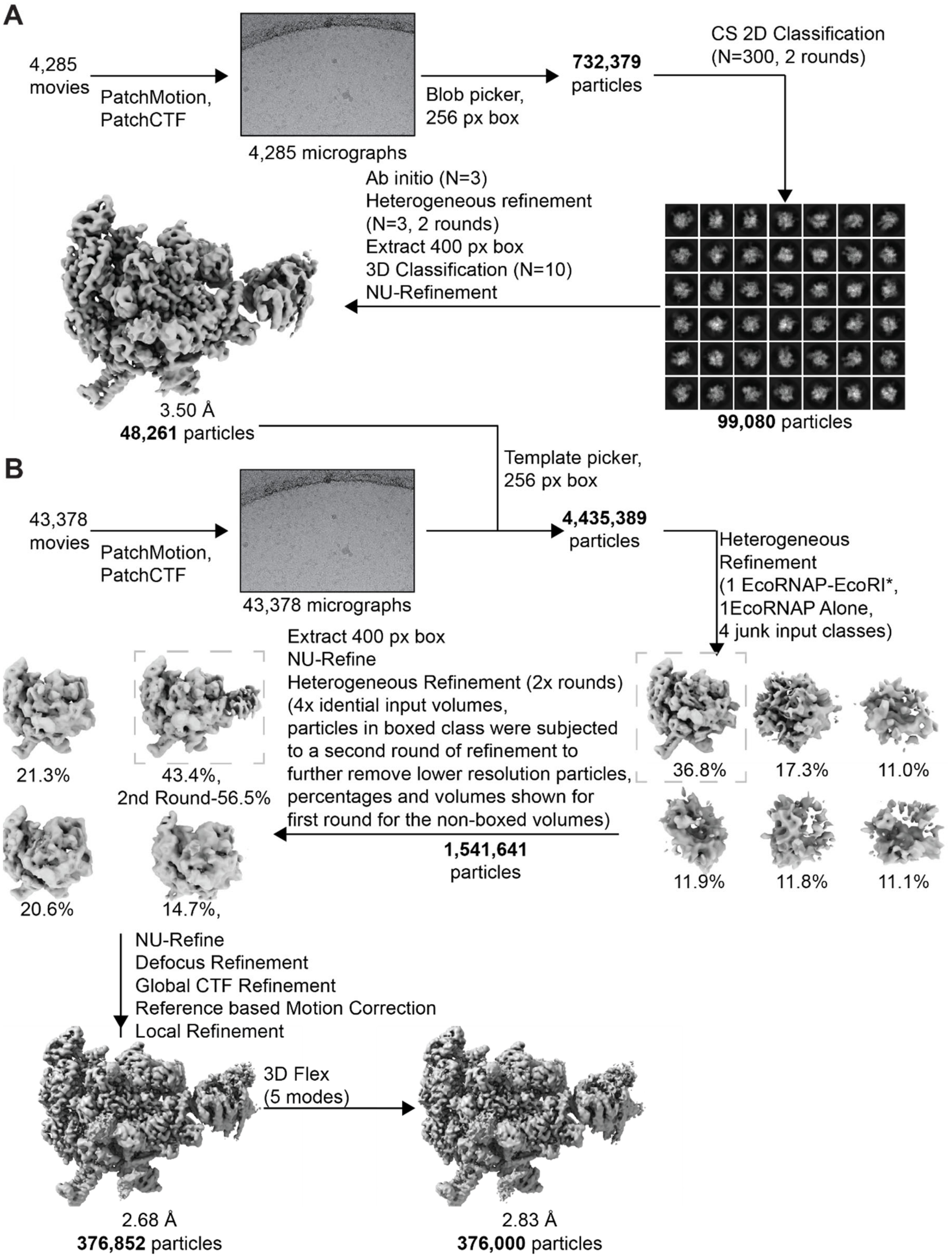


**Supplementary Figure 2. Cryo-EM processing of RNAP-EcoRI* collided complex.** Processing pipeline for the RNAP-EcoRI* complex for **[A]** a test dataset used to generate an initial reconstruction for template-based particle picking and **[B]** the full dataset.

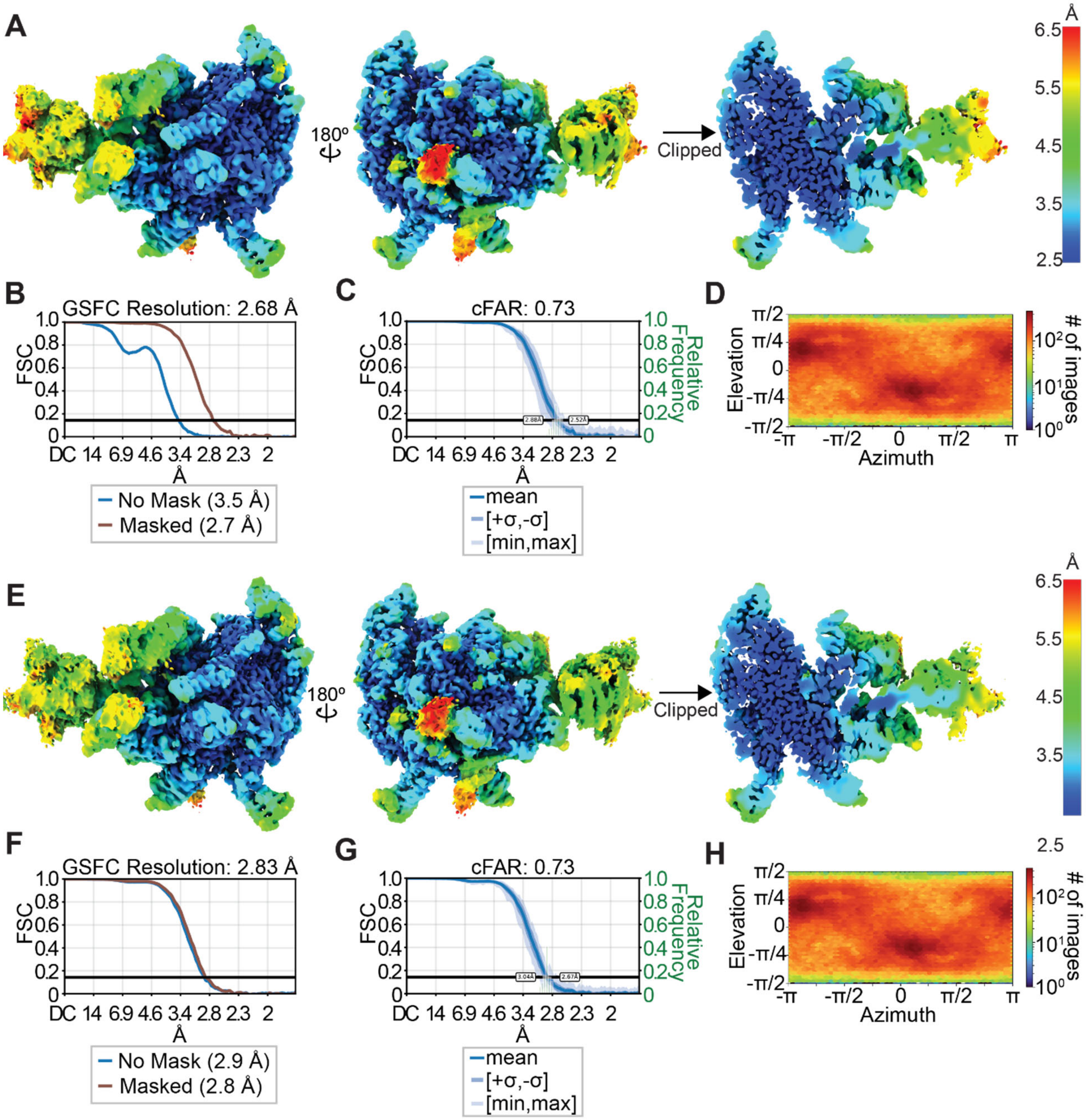


**Supplementary Figure 3. Resolution assessment and analysis of the RNAP-EcoRI* complex.**

**[A]** Local resolution estimation of non-uniform refinement consensus structure**,** as well as **[B]** global gold standard FSC curve, **[C]** conical FSC curve, and **[D]** viewing direction distribution.

**[E]** Local resolution estimation of 3DFlex refined consensus structure**,** as well as corresponding **[F]** global gold standard FSC curve, **[G]** conical FSC curve, and **[H]** viewing direction distribution.

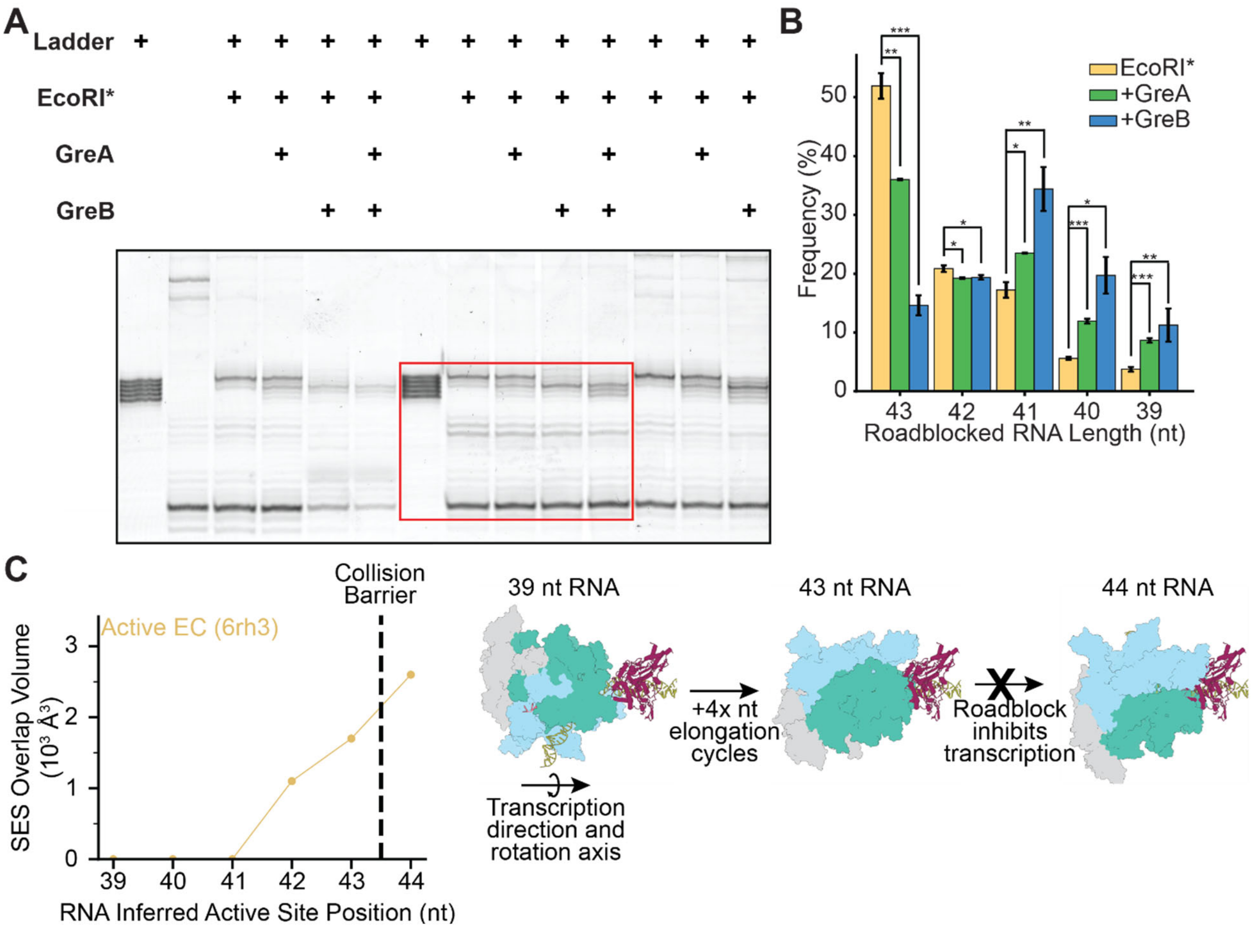


**Supplementary Figure 4. Additional analysis of the RNAP-EcoRI* collision complex.**

**[A]** Uncropped gel image of the backtracking assay shown in **Fig. 2B** (red box).

**[B]** Quantification of the assay presented in **A**. Band intensities were normalized for each lane. Data are presented as mean ± S.D. (N = 3). Conditions were compared via two-tailed, unpaired t test including Welch's correction. *$p < 0.05$; **$p < 0.01$; ***$p < 0.001$.

**[C]** Left: Quantification of overlapping volume enclosed by the solvent-excluded surfaces (SES) of the EcoRI* roadblock and RNAP as a function of position. Measurements were performed on an active EC (PDB 6RH3, representing a closed elongation complex) as the RNAP was computationally moved stepwise closer to the last position where nucleotide incorporation can occur (43 nt), then one position past the collision barrier (44 nt). Right: Snapshots at the indicated positions.

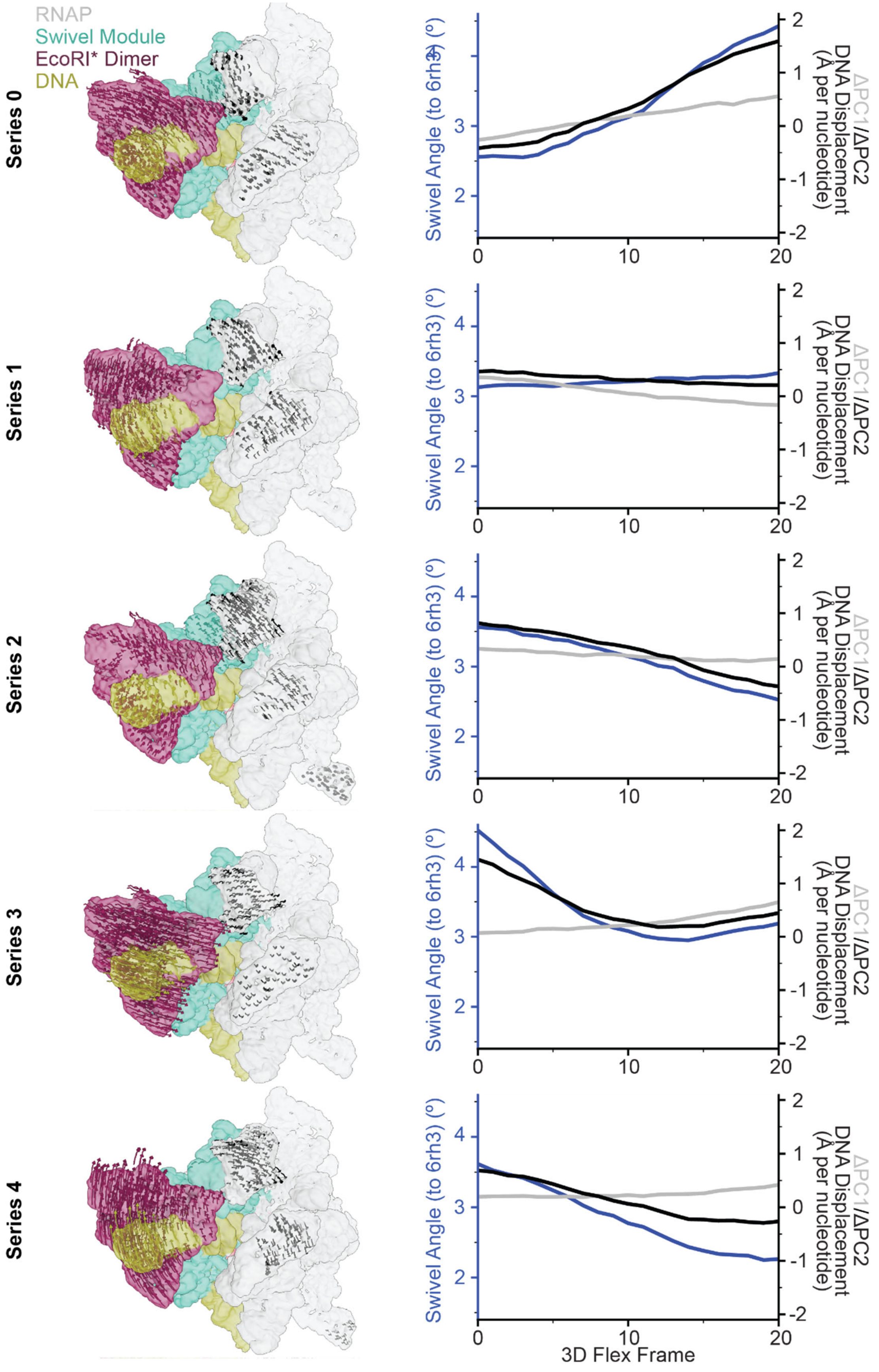
RNAP
Swivel Module
EcoRI* Dimer
DNA
Series 0
Series 1
Series 2
Series 3
Series 4
Swivel Angle (to 6rh3) (°)
ΔPC1/ΔPC2
DNA Displacement
(Å per nucleotide)
3D Flex Frame
0
10
20
2
3
4
-2
-1
0
1
2

**Supplementary Figure 5. Analysis of deformation-swivel coupling from 3DFlex trajectories for the EcoRI*-RNAP complex.** For the indicated 3DFlex trajectory series: (Left) Displacement vector plots for $C_\alpha$ atoms with >1.45 Å total RMSD change between first and last frame (arrows are magnified by 3 for display purposes). The vectors track the movement of the mobile residues at each frame of the 3DFlex trajectory. (Right) Plots of swivel angles and DNA displacements along the corresponding 3DFlex trajectories. Displacement values are calculated relative to the consensus reconstruction. Consensus class deformation in PC1/PC2 are set to zero.

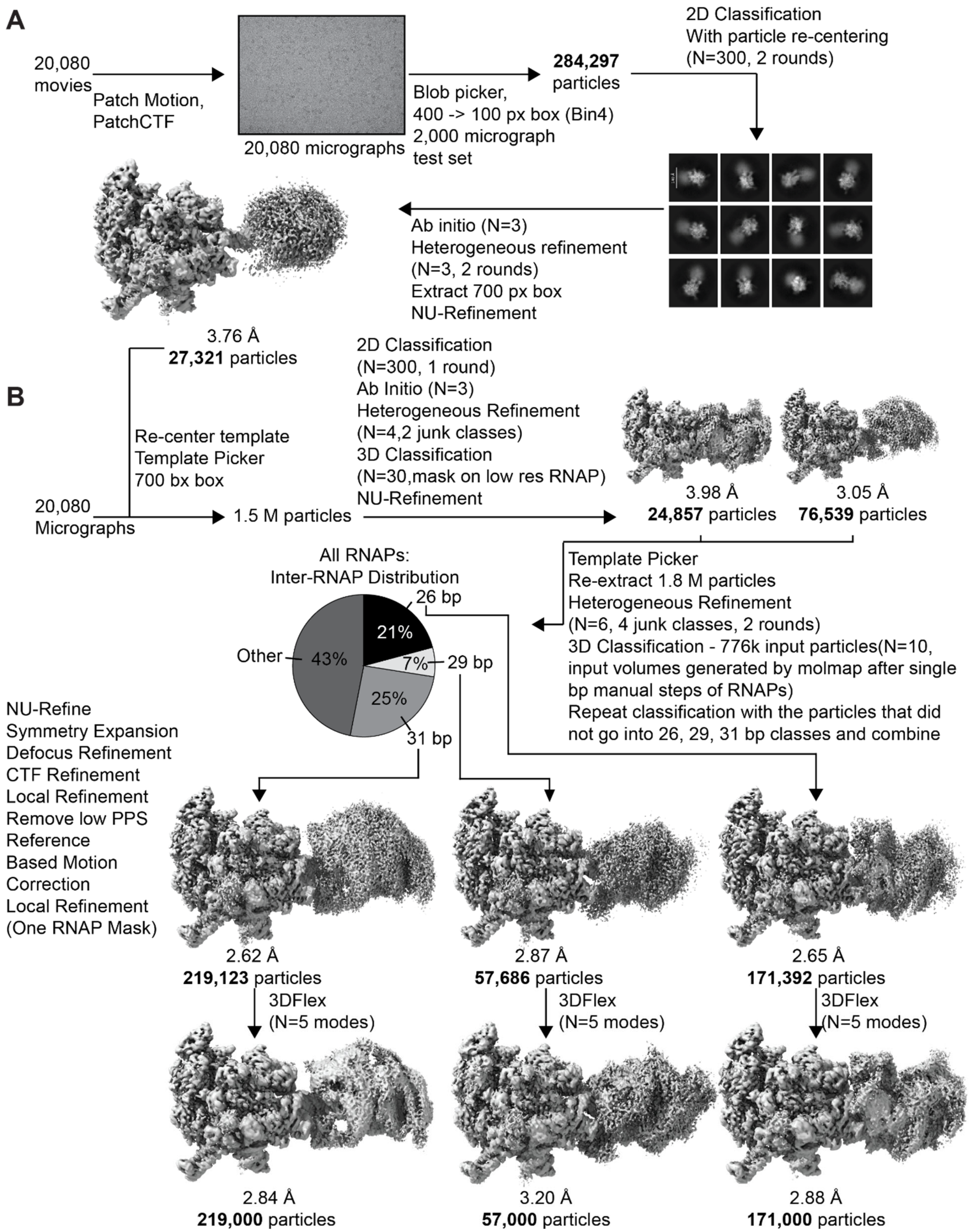


**Supplementary Figure 6. Cryo-EM processing of RNAP-RNAP head-on collision complex.**
Processing pipeline for the RNAP-RNAP complex for **[A]** a test dataset used to generate an initial reconstruction for template-based particle picking and **[B]** the full dataset.

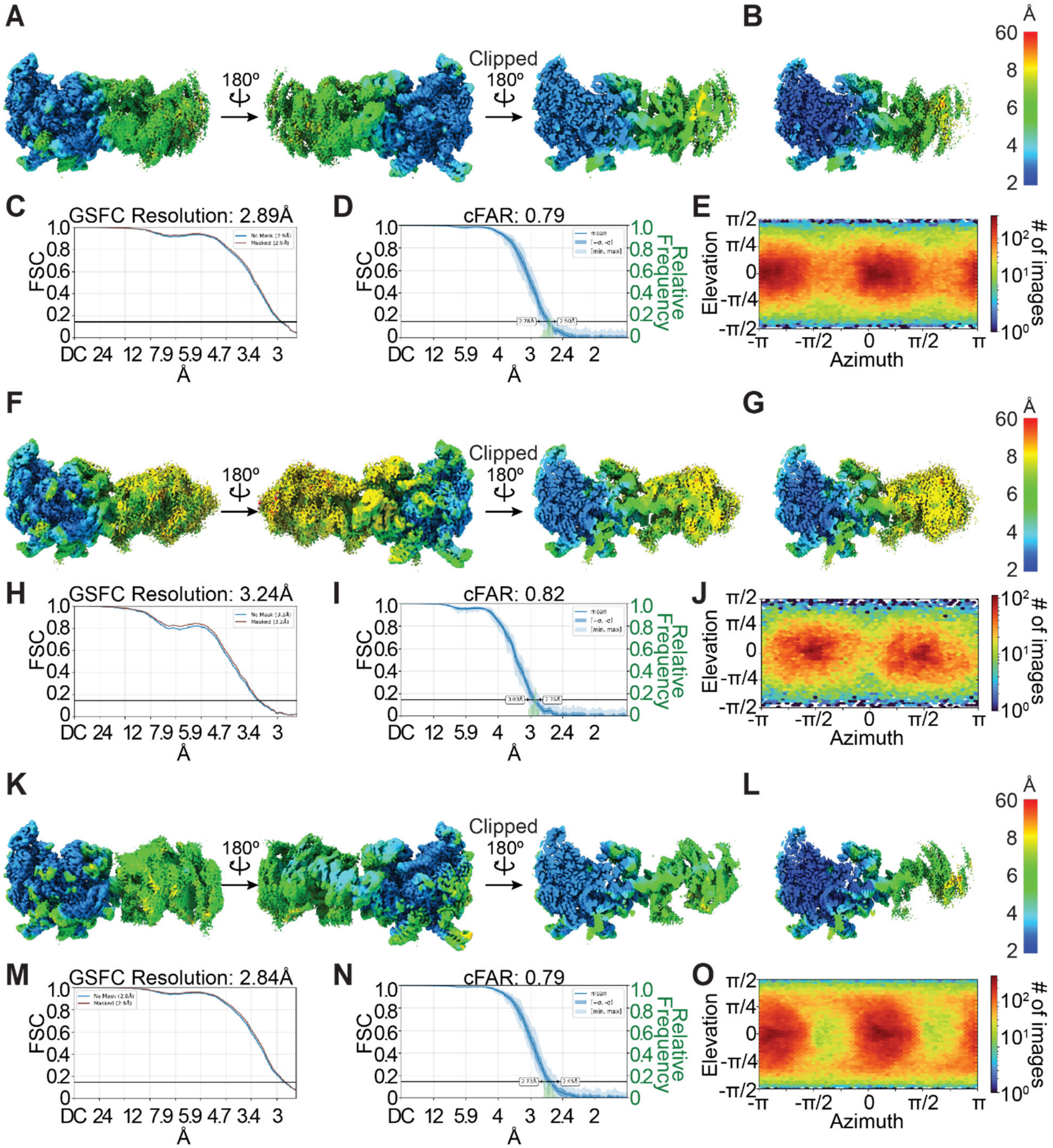


**Supplementary Figure 7. Resolution assessment and analysis of RNAP-RNAP complexes.**

Analysis of **[A-E]** 26-bp, **[F-J]** 29-bp, and **[K-O]** 31-bp inter-RNAP distance maps. **[A, F, K]** Local resolution of 3DFlex maps, including a clipped view of the center of the maps. **[B, G, L]** Clipped view of local resolution estimation of locally refined maps with a mask around the higher-resolution RNAP. **[C, H, M]** Global gold standard FSC curves for 3DFlex maps. **[D, I, N]** Conical FSC curves for the locally refined maps (masked around the higher-resolution RNAP). **[E, J, O]** Viewing direction distributions.

A

| | Total Reads | Initial Mapping | | TSS-Filtered | | Average Length(nt) | |
|---|---|---|---|---|---|---|---|
| | | Forward | Reverse | Forward | Reverse | Forward | Reverse |
| Hairpin | 50,360 | 16,687 | 16,187 | 5,408 | 4,081 | 75 | 80 |
| No hairpin | 51,976 | 14,872 | 19,619 | 6,470 | 6,828 | 50 | 62 |
| Hairpin pulldown | 12,076 | 4,366 | 2,696 | 1,789 | 608 | 125 | 104 |
| No hairpin pulldown | 89,104 | 30,297 | 26,409 | 8,075 | 5,945 | 93 | 88 |
| Reverse promoter only | 98,346 | 34,505 | 38,421 | 5,369 | 20,972 | — | 72 |
| Forward promoter only | 97,034 | 70,211 | 9,022 | 57,925 | 1,667 | 78 | — |

B

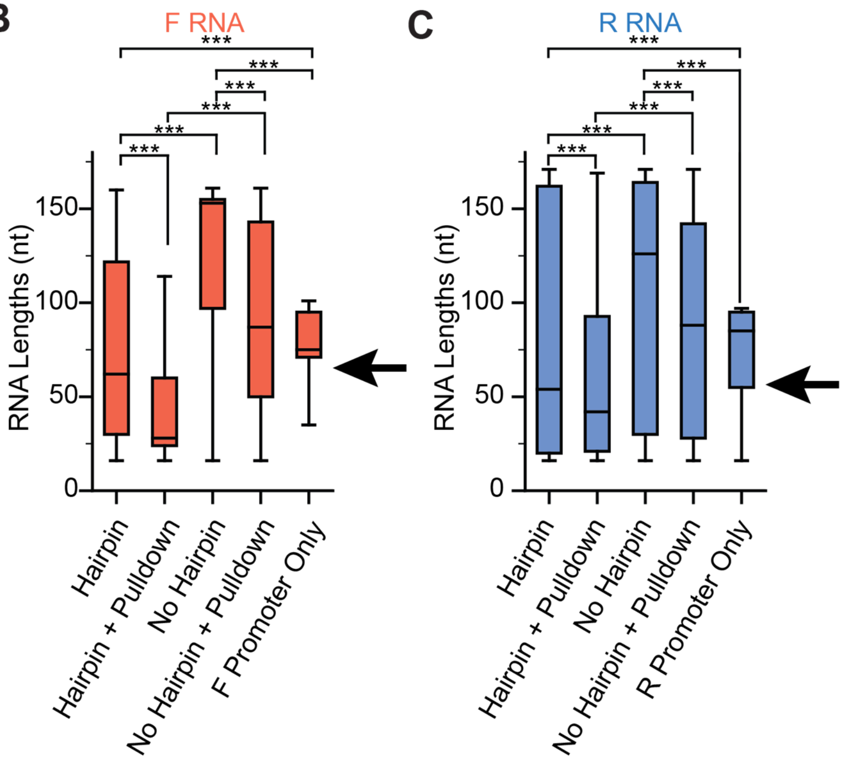


**Supplementary Figure 8. SEnd-seq data processing and filtering.**

**[A]** SEnd-seq read numbers, mapping, and filtering results for forward and reverse RNAs that have the correct transcription start site.

**[B, C]** Mapped, filtered RNA lengths under indicated conditions for **[B]** forward and **[C]** reverse RNAs. Arrows indicate the predicted RNA lengths if the hairpin was transcribed to completion. RNA products in the single promoter conditions were globally shorter due to the shorter template length. Conditions were compared by two-tailed, unpaired t-test including Welch’s correction and multiple-testing correction using the Benjamini-Hochberg method. ***$p < 0.001$.

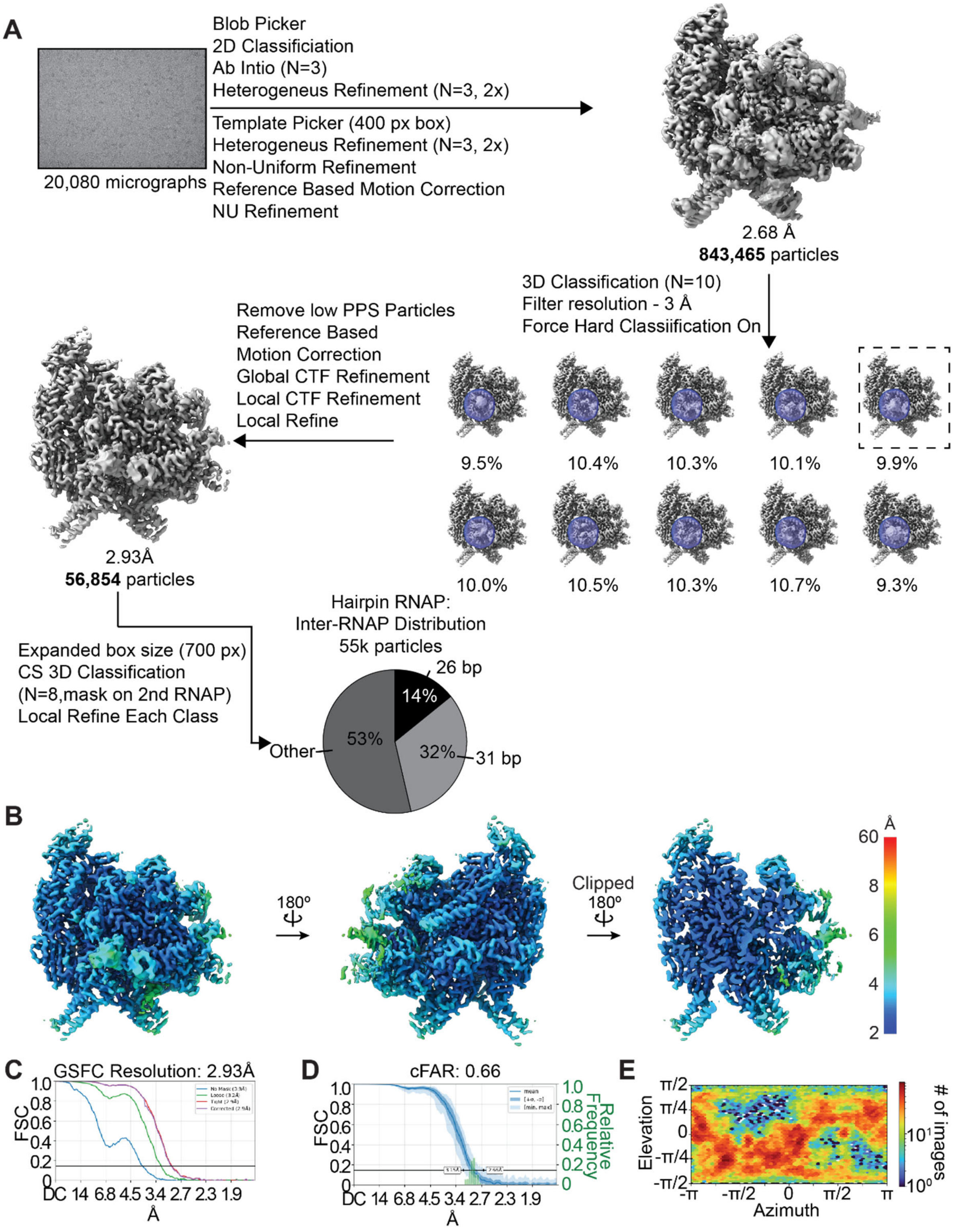


**Supplementary Figure 9. Cryo-EM processing and resolution assessment of RNAP featuring hairpin density.**

**[A]** Cryo-EM processing pipeline to enrich for hairpin-containing RNAPs.

**[B]** Local resolution estimation of locally refined hairpin-containing RNAP map.

**[C]** Global gold standard FSC curve.
**[D]** Conical FSC curve from the local refined map.
**[E]** Viewing direction distribution.

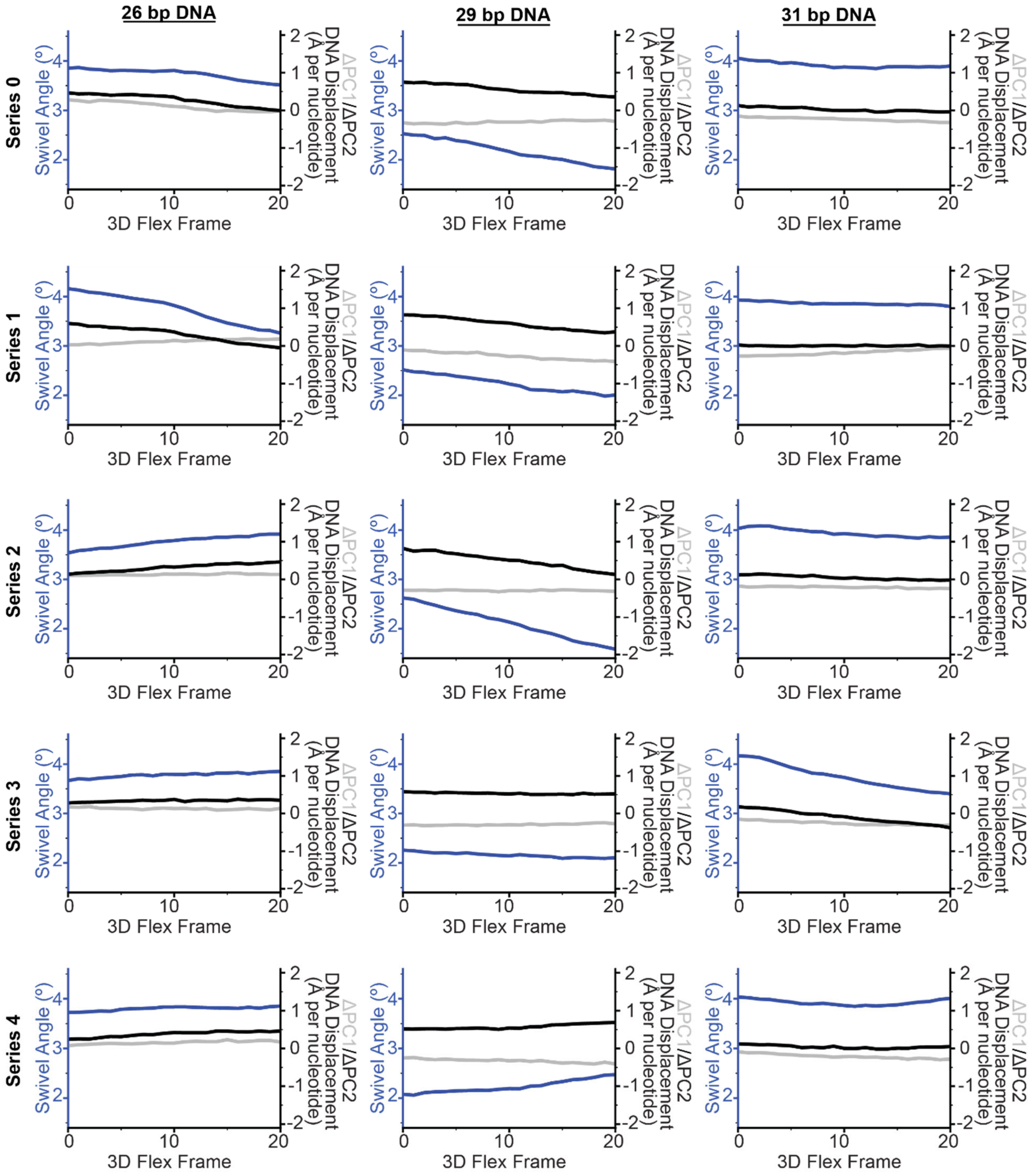


**Supplementary Figure 10. Analysis of DNA deformation / RNAP swiveling coupling in 3DFlex trajectories of RNAP-RNAP complexes.** Plots of RNAP swivel angle and DNA displacements along indicated 3DFlex trajectories for 26-bp (left), 29-bp (middle), and 31-bp (right) inter-RNAP distance complexes. Displacement values are calculated relative to the consensus reconstruction for each class. Consensus class deformation in PC1/PC2 are set to zero.

**Supplementary Movie 1.** (Left) 3DFlex variability trajectory movies for the first series for the RNAP-EcoRI* collision. Swivel module backbone trace is shown in blue, downstream DNA nucleotides analyzed shown in black, and the EcoRI dimer shown in magenta. Volume trajectories generated by 3DFlex are shown first and then made transparent to show the fit models aligned with the frame of reference used in this analysis. (Right) Animated plot showing the current swivel angle (top), PC1 displacement (middle), and PC2 displacement (bottom). Units for PC displacement are angstroms per nucleotide.

**Supplementary Movie 2.** Morph between the RNAP-EcoRI* collided complex and a superimposed EcoRI* protomer derived from the crystal structure (PDB 1ERI), highlighting rearrangement of the EcoRI* N-terminal helix (NTH) and repositioning of the N-terminal extension (NTE) toward the downstream DNA axis.

**Supplementary Movie 3.** (Left) 3DFlex variability trajectory movies for the first series for the head-on RNAP-RNAP collision (in order: the 26-bp, 29-bp and 31-bp inter-RNAP distance complexes). Swivel module backbone trace is shown in dark blue, downstream DNA nucleotides are colored matching the coloring in **Fig. 6**. Volume trajectories generated by 3DFlex are shown first and then made transparent to show the fit models aligned with the frame of reference used in this analysis. (Right) Animated plot showing the current swivel angle (top), PC1 displacement (middle), and PC2 displacement (bottom). Units for PC displacement are angstroms per nucleotide.

**Supplementary Table 1. Scaffold oligonucleotides and DNA sequences**

| **Description** | **Sequence (5'->3')**<br>Complementary region underlined when applicable<br>Oligonucleotides are DNA oligonucleotides unless specified others in the description | **Purity Notes** |
|---|---|---|
| *RNAP-EcoRI* Scaffold Oligos* | | |
| EcoRI* RNA Primer | Cy3 - UACUAUAGACACAAAGCCCAGCGG | PAGE (polyacrylamide gel electrophoresis)-purified. Ordered protected from Dharmacon |
| EcoRI* Scaffold NT | CTAGTCCATCAGTGTGCCCAGCGGTCAAATCTAGGTCTTATATATGTCACAATCGAGAATTCATTAGCTTACTTGG | PAGE-purified (IDT) |
| EcoRI* Scaffold T | CCAAGTAAGCTAATGAATTCTCGATTGTGACATATATAAGACCTAGATTTGACCGCTGGGCACACTGATGGACTAG | PAGE-purified (IDT) |
| Gre factor activity backtracked RNA 1 (15 nt) | AGCUAGA*G*G*UUUUUU | HPLC-purified (IDT)<br>* denotes phosphorothioate linkage |
| Gre factor activity backtracked RNA 1 (11 nt) | AGCUAGA*G*G*UU | HPLC-purified (IDT)<br>* denotes phosphorothioate linkage |
| Gre factor activity T | CTCTGAATCTCTTCCCCTCTAGCTTAGGACGTACTGACC | HPLC-purified (IDT) |
| Gre factor activity NT | GGTCAGTACGTCCATTCGATCTCCCGAAGAGATTCAGAG | HPLC-purified (IDT) |
| *RNAP-RNAP PCR Oligos*<br>5' biotin modifications were added to the primers to allow for streptavidin pulldown in certain experiments. Complementary region underlined. | | |
| T7A1 F Primer | CTGCCGCGACCAAGATTAATTTAAAATTTATCAAAAAGAGTATTGAC | No purification (IDT) |
| Lambda PR R Primer | GACGCCCTGGGGATAAATATCTAACACCGTGC | No purification (IDT) |
| No T7A1 F Primer | CTGCCGCGACTATGAGGAATTGTTATCCGCTCA | No purification (IDT) |
| No Lambda PR R Primer | GACGCCCTGGGTGTATTAAGGAGGTTGTATCCACACC | No purification (IDT) |
| | *Full PCR products (5'-3')*<br>Annotation: T7A1 (to +1) forward, Lambda PR (to +1) reverse, YnaJ-UspE hairpin, No Hairpin, **CTP deprivation block** | |

| | |
|---|---|
| T7A1-Lambda PR YnaJ-UspE Hairpin | CTGCCGCGACCAAGATTAATTTAAAATTTATCAAAAAGAGTATTGACTTAAAGTCTAACCTATAGGATACTTACAGCCATATGAGGAATTGTTAT**CC**GCTCACAATTCCAAAAAAATAGGCCCGATAACTCGGGCCTTGTCAGT**GG**ATACAACCTCCTTAATACATGCAACCATTATCACCGCCAGAGGTAAAATAGTCAACACGCACGGTGTTAGATATTTATCCCCAGGGCGTC |
| T7A1-Lambda PR No Hairpin | CTGCCGCGACCAAGATTAATTTAAAATTTATCAAAAAGAGTATTGACTTAAAGTCTAACCTATAGGATACTTACAGCCATATGAGGAATTGTTAT**CC**GCTCACAATTCCACCGATCATCAACTACATGAGAACATGCGAGGTGT**GG**ATACAACCTCCTTAATACATGCAACCATTATCACCGCCAGAGGTAAAATAGTCAACACGCACGGTGTTAGATATTTATCCCCAGGGCGTC |
| T7A1 Only YnaJ-UspE Hairpin | CTGCCGCGACCAAGATTAATTTAAAATTTATCAAAAAGAGTATTGACTTAAAGTCTAACCTATAGGATACTTACAGCCATATGAGGAATTGTTAT**CC**GCTCACAATTCCAAAAAAATAGGCCCGATAACTCGGGCCTTGTCAGTGGATACAACCTCCTTAATACACCCAGGGCGTC |
| LambdaPR Only YnaJ-UspE Hairpin | CTGCCGCGACTATGAGGAATTGTTATCCGCTCACAATTCCAAAAAAATAGGCCCGATAACTCGGGCCTTGTCAGT**GG**ATACAACCTCCTTAATACATGCAACCATTATCACCGCCAGAGGTAAAATAGTCAACACGCACGGTGTTAGATATTTATCCCCAGGGCGTC |

### Supplementary Table 2. Cryo-EM data collection, refinement, and validation statistics

| | RNAP-EcoRI* Collision | Head-on RNAP-RNAP Collision: 26 bp distance | Head-on RNAP-RNAP Collision: 29 bp distance | Head-on RNAP-RNAP Collision: 31 bp distance | Head-on RNAP-RNAP Collision: RNA Hairpin |
|---|---|---|---|---|---|
| PDB ID | XXXX | XXXX | XXXX | XXXX | XXXX |
| EMDB | YYYYY | YYYYY | YYYYY | YYYYY | YYYYY |
| **Data collection and processing** | | | | | |
| Microscope | Titan Krios | Titan Krios | | | |
| Voltage (kV) | 300 | 300 | | | |
| Detector | Gatan K3 | Gatan K3 | | | |
| Magnification | 81,000 | 105,000 | | | |
| Electron Exposure (e-Å$^{-2}$) | 47.3 | 48.8 | | | |
| Exposure rate (e-/pixel/s) | 25 | 25 | | | |
| Calibrated pixel size (Å) | 0.86 | 0.847 | | | |
| Defocus Range (μm) | -0.8 to -2.0 | -0.8 to -2.0 | | | |
| Micrographs (no.) | 43,378 | 20,080 | | | |
| Initial particle images (no.) | 4,435,389 | 1,801,942 | | | 2,946,928 |
| Symmetry applied | C1 | C1 | C1 | C1 | C1 |
| Symmetry Expanded? | No | Yes, pseudo C2 | Yes, pseudo C2 | Yes, pseudo C2 | No |
| Final particle images (no.) | 376,852 | 219,123 | 57,686 | 171,392 | 56,854 |
| Local refined map resolution (Å) | 2.68 | 2.65 | 2.87 | 2.62 | 2.93 |
| 3DFlex refined map resolution (Å) | 2.83 | 2.88 | 3.20 | 2.84 | - |
| FSC threshold | 0.143 | 0.143 | 0.143 | 0.143 | 0.143 |
| **Refinement** | | | | | |
| Initial model(s) used (PDB Code) | 6RIP,1ERI | 6RIP | 6RIP | 6RIP | 6RIP, 6ASX |
| Model Composition | 1 RNAP, 2 EcoRI | 1 RNAP | 1 RNAP | 1 RNAP | 1 RNAP |
| Non-hydrogen atoms | 31,469 | 26,434 | 26,422 | 26,434 | 26,653 |
| Protein Residues | 3,729 | 3,194 | 3,197 | 3,194 | 3,194 |
| RNA bases | 13 | 13 | 13 | 13 | 25 |
| DNA bases | 96 | 56 | 54 | 56 | 54 |
| Ligands ($Zn^{2+}$/$Mg^{2+}$) | 2/1 | 2/1 | 2/1 | 2/1 | 2/1 |
| Map sharpening B factor (Å$^2$) | -91.22 | -66.73 | -56.27 | -68.32 | -65.90 |
| Average B-factor protein (Å$^2$) | 71.59 | 53.34 | 70.61 | 29.54 | 63.77 |
| Average B-factor nucleotide (Å$^2$) | 39.31 | 42.08 | 41.81 | 44.64 | 53.67 |
| **RMS deviations** | | | | | |
| Bond lengths (Å) | 0.004 | 0.004 | 0.004 | 0.005 | 0.004 |
| Bond angles (°) | 0.916 | 0.972 | 0.944 | 0.967 | 0.945 |
| **Ramachandran plot** | | | | | |
| Favored (%) | 98.49 | 98.27 | 97.67 | 98.36 | 97.99 |
| Allowed (%) | 1.51 | 1.73 | 2.33 | 1.64 | 2.01 |
| Outliers (%) | 0.00 | 0.00 | 0.00 | 0.00 | 0.00 |
| **Validation** | | | | | |
| Molprobity score | 0.77 | 0.69 | 0.77 | 0.61 | 0.77 |
| Clash score | 0.88 | 0.58 | 0.58 | 0.31 | 0.89 |
| Rotamer outliers (%) | 0.00 | 0.00 | 0.04 | 0.00 | 0.00 |

**Supplementary Table 3. Domains defined for rigid body fitting**

| Description | Chain ID and Residue Selection |
|---|---|
| *For high-resolution RNAP models: protein only* | |
| α Protomer 1 NTD Part 1 | /A:53-178 |
| α Protomer 1 NTD Part 2 | /A:1-52,179-236 |
| α Protomer 1 NTD Part 1 | /B:53-178 |
| α Protomer 1 NTD Part 2 | /B:1-52,179-234 |
| Structural core | /C:2-27,147-152,445-455,520-713,786-828,1060-1318<br>/D:343-368,421-644,705-786,1345-1376,1503 |
| β protrusion | /C:28-146,456-519 |
| β lobe | /C:153-225,340-444 |
| B SI1 | /C:226-339 |
| Rim helices | /C:645-704 |
| B stalk | /C:714-785 |
| β flap | /C:829-894,910-937,1040-1059 |
| β SI2 | /C:938-1039 |
| Clamp | /C:1319-1342/D:1-342,1318-1344/D:1501 |
| β' sw1 | /D:369-420 |
| Shelf | /D:787-935,1135-1150,1216-1317,1502 |
| β' SI3 Part 1 | /D:948-1023 |
| β' SI3 Part 2 | /D:1024-1126 |
| Jaw | /D:1151-1215 |
| ω | /E |
| *Nucleic acids and other protein components* | |
| RNAP-EcoRI* | |
| Upstream DNA and RNA | /N:16-50/T:27-61/R |
| Downstream DNA and EcoRI* | /F/G/N:51-76/T:1-26 |
| Head-on RNAP-RNAP Collision: 26 bp distance | |
| Upstream DNA | /N:7-35/T:39-66/R |
| Downstream DNA and other RNAP | /F-J/N:36-67/T:25-38/S |
| Head-on RNAP-RNAP Collision: 29 bp distance | |
| Upstream DNA | /N:11-37/T:40-65/R |
| Downstream DNA and other RNAP | /F-J/N:38-66/T:11-39/S |
| Head-on RNAP-RNAP Collision: 31 bp distance | |
| Upstream DNA | /N:11-40/T:39-67/R |
| Downstream DNA and other RNAP | /F-J/N:41-68/T:11-38/S |
